# Transcriptomic differences underlying fin sexual dichromatism and male color polymorphism in bluefin killifish (*Lucania goodei*)

**DOI:** 10.64898/2026.09.04.749336

**Authors:** Ratna Karatgi, Chi-Hing Christina Cheng, Giovanni Madrigal, Julian M Catchen, Rebecca C. Fuller

## Abstract

Color polymorphisms offer a powerful lens for examining how selection shapes genetic and phenotypic variation. Genomic analyses of color variation provide deeper insight into how genetic differences and phenotypic plasticity contribute to trait variation. Here, we use the bluefin killifish (*Lucania goodei*) to study this relationship. Male fins can display either red or yellow coloration based on pigments (pterins) or blue structural coloration (iridophores) often induced in UV-depauperate environments. Female fins lack pigmentation. To characterize the molecular basis of this variation, we assembled a high-quality genome and examined gene expression in the anal fins of male and female bluefin killifish to identify the molecular mechanisms underlying color variation. Our gene expression analysis found sex differences driven by male-biased upregulation of melanin synthesis and canonical pigment pathway genes. We additionally identified that the progesterone sex receptor was upregulated in females. Among male color morphs, several genes involved in xanthophore and iridophore formation differed between blue and non-blue males, including the transcription factor *tfec*, and iridophore-patterning genes *atic*, *pnp4a*, and *sox10*. Extraocular opsins, which are involved in non-image-forming light perception, were expressed at low levels across all fins, but showed population- and morph-specific patterns; notably, *rgrb* (retinal G protein coupled receptor b) was upregulated in blue males. To the best of our knowledge, this is the first report of an extraocular opsin varying as a function of population and color morph. Together, these results identify the pigment pathways underlying male-limited color polymorphisms and reveal candidate genes likely involved in light-sensitive phenotypic plasticity.

## Introduction

Trait variation is ubiquitous across organisms; color polymorphisms, which are discrete variations in color-based traits, are some of the most easily observable forms of phenotypic variation (Endler, 1978; Ford, 1945; McKinnon & Pierotti, 2010). Color polymorphisms were the subject of some of the earliest studies demonstrating how selection acts on trait variation in species, furthering our understanding of the roles of frequency dependence, spatial and temporal variation in selection, and predation on trait evolution (Ford, 1975; Giesel, 1971; Jameson & Pequegnat, 1971; Halkka et al., 1980; Hagen et al., 1980; Hairston, 1981; Endler, 1986; Hoekstra, 2006; McKinnon & Pierotti, 2010; Wellenreuther et al., 2014; Orteu & Jiggins, 2020). Additionally, color polymorphisms that are sexually dimorphic have improved our understanding of sexual selection through crucial intraspecific factors such as mate choice, competition, and sexual antagonism, and the interaction of sexual and natural selection in shaping traits (Dijkstra et al., 2007; Gray & McKinnon, 2007; Mitchem et al., 2018). Careful breeding studies have often provided insights into the genetic and environmental nature of color polymorphisms that were foundational to our early understanding of evolutionary biology (Allen, 1904; Ford, 1945; Hoekstra, 2006; Morgan, 1914a, 1914b; Pool & Aquadro, 2007; Wellenreuther et al., 2014). Yet, despite the importance and functional significance of color variation, we often lack a complete understanding of the underlying mechanisms generating these variable phenotypes.

Understanding the genetic and physiological changes responsible for variation in coloration provides insight into the proximate mechanisms of trait variation. In particular, studying the gene expression differences leading to polymorphic color traits can help us understand the cellular and molecular mechanisms underlying specific traits, indicate candidate loci for further genomic analysis, and potentially identify regulatory differences (Ahi et al., 2020; Ahi & Sefc, 2017; Chauhan et al., 2016; Darolti & Mank, 2023; Djurdjevič et al., 2019; Henning et al., 2013; Ng’oma et al., 2014; Recknagel et al., 2024; Zhang et al., 2017). This can also be especially helpful for studying color-based traits that are plastic responses to environmental changes (Henning et al., 2013; Lafuente & Beldade, 2019). In this study, we examine gene expression in the bluefin killifish system to describe variation in coloration at multiple levels: (1) between males and females, (2) among different color morphs, and (3) among populations and lighting environments (Figure 1).

**Figure 1.**
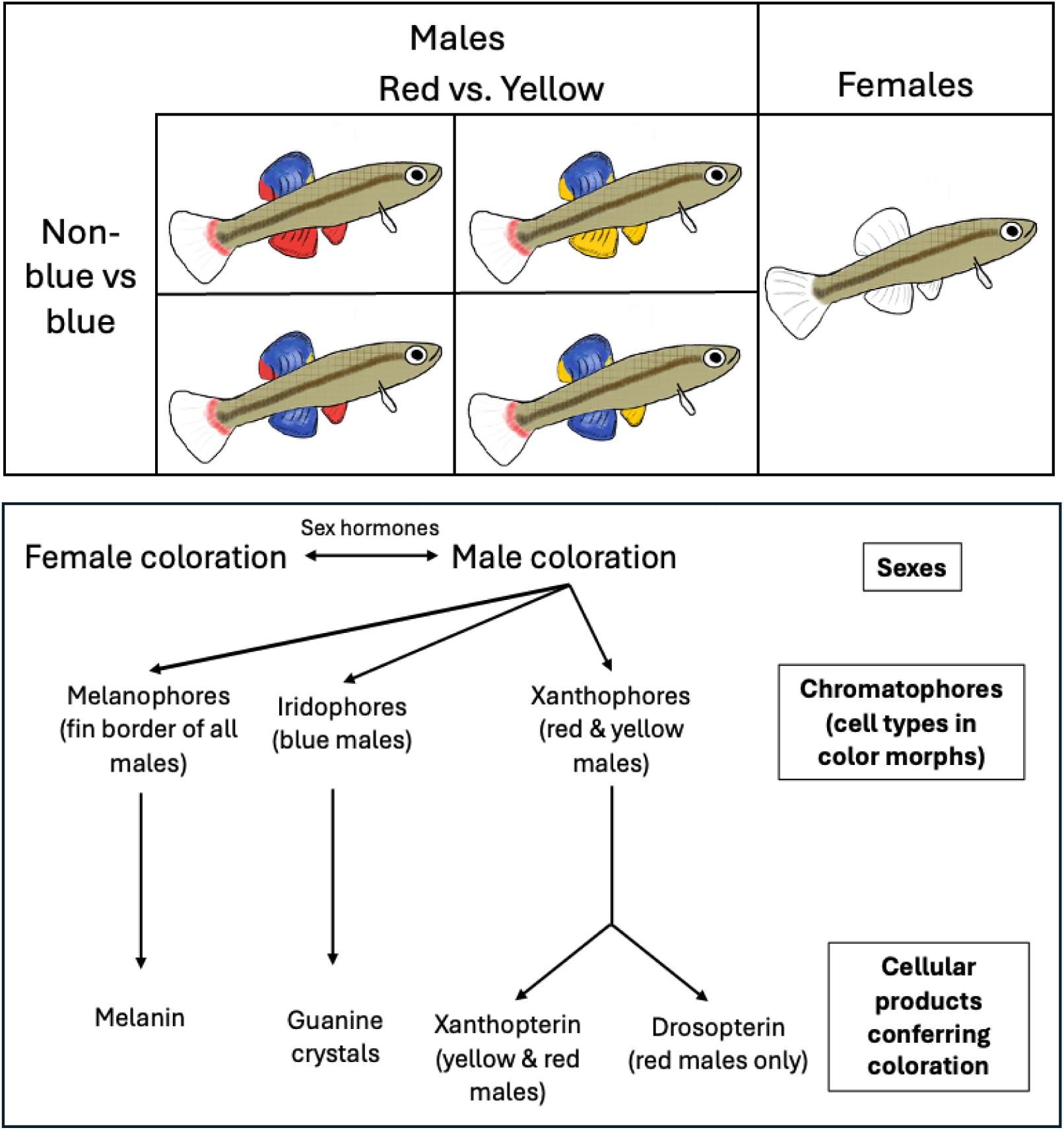
(top) Male and female bluefin killifish. Males can have red or yellow anal fins, or suppress this axis of coloration and exhibit blue anal fins. Males with red or yellow anal fins express red or yellow, respectively, on their pelvic fins. However, males with blue anal fins typically express either red or yellow pelvic fins. Females lack fin coloration. (bottom) Schematic for cell and pigment differences in bluefin killifish. We expect that sex steroid hormones underlie color differences between males and females. All males possess some amount of black melanin pigmentation on their fins, presumably due to melanin pigment contained in melanophores. Blue fins lack pigmentation, and their coloration is instead due to structural coloration arising from the organization of guanine crystals and reflective platelets within iridophore cells. Red and yellow coloration is due to pteridine pigments within xanthophores. Anal fins with red coloration have both xanthopterin and drosopterin pigments, whereas yellow fins only contain xanthopterin.

The first major axis of color variation in bluefin killifish (Figure 1) is between the sexes. In bluefin killifish, females have colorless anal fins, whereas male anal fins have a black melanin border and either pterin pigments (red or yellow) or structural coloration (blue) (Johnson & Fuller, 2015). Comparisons of sex differences in gene expression provide insights into which specific molecular pathways are involved in producing sexually dimorphic coloration and how this varies with different color morphs. In addition, sexually dimorphic traits often arise from differential gene regulation between males and females, yet the regulatory mechanisms generating these expression differences are not always fully characterized across systems (Ellegren & Parsch, 2007; Ingleby et al., 2015; Pointer et al., 2013). Steroid hormone-mediated signaling has long been implicated in shaping sex- and morph-specific gene expression.

Although steroid hormones are not themselves gene products, the enzymes involved in their synthesis and their receptors are; thus, examining expression patterns of hormone receptors offers a promising approach for identifying candidate regulatory pathways linking endocrine signals to sexual dimorphism and color polymorphism (Forlano et al., 2010; Mank, 2023; Ogino et al., 2023; Prazdnikov, 2022, 2025; Schuppe et al., 2017). In bluefin killifish, multiple lines of evidence suggest that coloration is hormonally regulated: juveniles resemble adult females (Foster, 1967), and experimental androgen treatment of females induces the expression of male-typical color patterns (Fuller & Travis 2004), suggesting that females are sensitive to androgens.

Beyond sexual dimorphism, bluefin killifish show striking male color variation along two axes: (1) presence or absence of blue structural color, and (2) the red–yellow pterin pigment axis (Figure 1). Sexually mature males display solid red, yellow, or blue anal fins, and the anal fin coloration can be correlated with that of pelvic and posterior dorsal fins. Red and yellow males have matching pelvic and posterior dorsal fins, whereas blue males retain red or yellow hues on those fins. Red and yellow colors arise from the pteridine pigments, drosopterin and xanthopterin, respectively, with red males expressing both drosopterin and xanthopterin and yellow males expressing only xanthopterin (Johnson & Fuller, 2015). Blue coloration instead results from the presence of guanine crystal-containing iridophore cells that reflect light structurally. Thus, blue male anal fins are characterized by suppression of pterin expression and the production of reflective intracellular structures, offering an ideal system for dissecting pigment-based and structural coloration pathways.

Much of what we know about how such colors are produced comes from research on chromatophores—the pigment-containing cells responsible for coloration in fish, amphibians, and reptiles (Bagnara et al., 1979; Parichy, 2006a; Andrade & Carneiro, 2021; Hashimoto et al., 2021). Studies in model systems such as zebrafish, cichlids, and medaka have identified the chromatophore types, enzymatic pathways, and transcription factors underlying pigment synthesis and cell differentiation (Lister, 2002; Maan & Sefc, 2013; Parichy, 2006a; Schartl et al., 2016; Patterson & Parichy, 2019). Bluefin killifish share many of these pigment cell types— melanophores (black melanin), xanthophores (red or yellow pterins), and iridophores. However, the genetic mechanisms driving color variation (especially non-carotenoid-based colors) in wild, non-model species like the bluefin killifish remain largely unexplored.

Finally, there is variation in coloration in bluefin killifish that is attributable to the lighting environment. The red versus yellow axis is largely genetic, with alleles at a locus of large effect influencing male coloration (Fuller & Travis, 2004; Fuller et al. 2022). Blue coloration is controlled by a combination of genetic variation, phenotypic plasticity, and genetic variation in phenotypic plasticity (Fuller et al., 2022). Three separate studies have shown that swamp population males originating from tannin-stained rivers are more likely to express blue coloration when reared in tea-stained environments mimicking swamp rivers (Fuller et al., 2022; Fuller & Travis, 2004; Karatgi, 2026). However, this plasticity is itself variable within and among populations. Comparisons between fish from a spring population, where the habitat exhibits a clear lighting environment, and a swamp population showed greater plasticity in the swamp population than the spring population, indicating that there is genetic variation in the plasticity in the expression of the blue phenotype (Fuller et al., 2022). Plastic traits often have complex genetic architectures, and analyzing gene expression differences between alternative plastic phenotypes is key to uncovering the mechanisms that drive plasticity (Henning et al., 2013; Hosseini et al., 2019; Lafuente & Beldade, 2019).

In this study, we use RNA sequencing to examine the gene expression differences underlying sexual dichromatism and male-limited color polymorphism in bluefin killifish. We ask the following questions: (a) What genes underlie sex differences in fin coloration? (b) Are all male morphs equally transcriptionally dissimilar to females? (c) What are the genes underlying the pterin pigment-based red- and yellow-colored fins vs the structural iridophore-based, blue-colored fins? To what extent are these differences conserved between populations? (d) Are there candidate genes that might contribute to light sensitivity in blue morphs? To address these questions, we used pairwise contrasts to test for differential gene expression among the experimental groups and complemented these genome-wide analyses with targeted examinations of *a priori* candidate genes with roles in pigmentation, sex differences, and photoreception. For the candidate gene analyses, we examined the following: (a) to identify the potential hormonal mechanisms underlying sexual dichromatism, we examined whether sex steroid receptor genes differed as a function of sex. (b) Among male morphs, we examined the effect of blue coloration (blue vs. non-blue) on the expression of thyroid hormone receptor genes, as they are known to be involved in regulating the differentiation of pigment-containing chromatophores. Additionally, we examined whether genes known to be involved in the pteridine synthesis pathway leading to red and yellow pterin coloration and genes involved in iridophore specification differed as a function of male blue status (presence/absence of blue coloration). (d) Finally, we examined whether non-visual opsin genes, which are involved in non-image-forming photoreception and known to be expressed in fin tissues, differed as a function of population and male morph type.

## Methods

### Genome Assembly

One female bluefin killifish was collected in the Wakulla River, Wakulla County, FL, in January 2021 and returned to the University of Illinois, Urbana-Champaign. The fish was euthanized, and DNA was extracted immediately using tissue from the entire set of gills. High Molecular Weight (HMW) DNA was extracted using the Nanobind Tissue Big DNA kit (Circulomics/PacBio Sciences), yielding ∼35 µg. DNA was assessed for purity with a Nanodrop One (Fisher Scientific), concentration by fluorometry with a Qubit v.3 (Invitrogen), and integrity and size on a 1% agarose gel. The HMW DNA was submitted to the Roy J. Carver Biotechnology Center at the University of Illinois Urbana-Champaign for long-read HiFi sequencing of one SMRT cell on a PacBio Sequel II instrument. The DNA was sheared with Megaruptor (Diagenode) to target insert lengths and converted into a library using the SMRTBell Express Template Prep kit 2.0 (PacBio). The library was sequenced on one SMRT cell 8M on a PacBio Sequel II for 30 hrs of data collection, and circular consensus sequence (CCS) calls were performed using SMRTLink v.9.0. The sequencing generated 1.98 million reads with a mean read length of 11.4 Kilobases (Kb), totaling 22.56 Gigabases (Gb) of read data. DNA from the same individual was also extracted and processed into the Omni-C variant of a chromosomal conformation capture library and submitted for sequencing on an Illumina NovaSeq 6000 instrument generating 2×150bp reads. The genome was assembled into contigs using HiFiasm version 0.19.8 (Cheng et al., 2021), using default parameters, and supplying the PacBio long-reads, and the Omni-C paired-end short-read data. In addition, a different fish was collected from the Santa Fe River near Rum Island Springs County Park (Columbia, Co., FL) in 2016. Its DNA was extracted and submitted for 10X Genomics sequencing on an Illumina HiSeq 4000 instrument, generating 2×150bp linked reads in 2017. The 10X Genomics linked reads were aligned to the resulting contigs using BWA version 0.7.17-r1188 (Li, 2013) and were scaffolded with Arcs version 1.2.7 (Yeo et al., 2018) and LINKS version 2.0.1 (Warren et al., 2015). A second round of scaffolding was then performed by first aligning the Omni-C paired-end short-reads with BWA to the Arcs-scaffolded data and then further scaffolding with YAHS version 1.2 (Zhou et al., 2023).

The resulting genome was annotated using the BRAKER3 pipeline (Gabriel et al., 2024), first identifying repeats using RepeatModeler version 2.0.4 (Flynn et al., 2020) and then using the generated repeat library to mask the genome with RepeatMasker version 4.1.5 (Smit et al., 2015). We aligned paired-end short-read RNA data to the genome using HiSat2 version 2.2.1 (Kim et al., 2019) and then executed the BRAKER3 pipeline. We supplied BRAKER3 with proteins from OrthoDB version 11(Kuznetsov et al., 2023) and from the *Fundulus heteroclitus* reference genome (NCBI accession GCF_011125445.2), which then internally ran Augustus and GeneMark to annotate based on proteins and RNA reads, respectively, with the best transcripts being selected by the internal TSEBRA program (Gabriel et al., 2021). To link gene models to known gene functions, we executed Interproscan version 5.68 (Quevillon et al., 2005), and we used Synolog (Madrigal & Catchen, 2026) to link genes to common gene names based on the zebrafish reference genome (GRCz11, NCBI accession GCA_000002035.4).

Briefly, Synolog identifies orthologs through an initial reciprocal best hit method and then applies a synteny-based approach to assign orthologs to unmatched genes based on shared gene neighborhoods and sequence similarity. Finally, we renamed the resulting assembled chromosomes using Synolog based on conserved synteny with the *Xiphophorus maculatus* reference genome (NCBI accession GCF_002775205.1).

### Fish Collection, Animal Husbandry, Tissue Collection and RNA Extraction

Male and female bluefin killifish were caught using seine nets from two different populations, Rainbow River (a spring population), and the Everglades (a swamp population), in Florida in April 2023. Fish were transported to a greenhouse at the University of Illinois Urbana-Champaign, where they were initially housed in 400-liter stock tanks with large sponge filters that kept the water column aerated and free of nitrogenous wastes. Fish were fed daily with frozen brine shrimp and supplemented with aquatic invertebrates (Daphnia) and algae that naturally grow in the stock tanks. The Everglades population was housed in tea-stained water, mimicking their tannin-stained swamp habitat. Tea-stained water was created by adding instant, decaffeinated, no-sugar, no-lemon tea powder to the water. Rainbow River fish were housed in clear water. Hence, the fish were housed in lighting environments that were similar to their native habitats. One month before tissue collection, male-female pairs were moved to 110-liter tanks and held until tissue harvesting. Housing a male with females increases the brightness and saturation of male anal fin color (Johnson & Fuller, 2015), and this was done to ensure that an adequate amount of RNA could be extracted from fin tissues.

We extracted RNA from anal fin tissues from females and male morphs from both populations. The male morph type includes males with red anal fins (red morph), males with yellow anal fins (yellow morph), males with blue anal fins and red pelvic fins (blue-red morph), and males with blue anal fins and yellow pelvic fins (blue-yellow morph). We used four replicate individuals from each population for each of the male morphs and the females, except for the blue-red morph from the Rainbow River population, where we had three replicates. In total, fin tissue was collected from 39 individual fish. Tissue was collected between 10 am-1 pm daily to minimize potential gene expression variation due to diurnal rhythms.

Fish were euthanized with an overdose of buffered Syncaine MS-222 (Fisher Scientific). The anal fin was promptly dissected and removed from the fish. Tissue samples were stored in RNAlater (Invitrogen) at 4°C, and extraction was done within a week of dissection. Tissues were homogenized in Trizol reagent (Invitrogen) with 0.5mm zirconium oxide beads using a Bullet Blender (Next Advance). Total RNA was isolated using Trizol reagent as per the manufacturer’s recommendations. Samples were treated with DNAse I (New England Biolabs) to remove genomic DNA, and RNA was precipitated with sodium acetate. RNA quality and quantity were assessed using a Nanodrop spectrophotometer and Qubit 2.0 Fluorometer. Samples were immediately stored at -20°C and submitted for sequencing within one month of extraction.

### RNA sequencing

RNA sequencing libraries were prepared with the Kapa Hyper Stranded mRNA library kit (Roche). The libraries were pooled, quantified by qPCR, and sequenced on two 10B lanes for 101 cycles from one end of the fragments on a NovaSeq X Plus with V1.0 sequencing kits.

FASTQ files were generated and demultiplexed with the bcl2fastq v2.20 Conversion Software (Illumina). Library preparation and sequencing were carried out at the Roy J. Carver Biotechnology Center at the University of Illinois Urbana-Champaign.

An average of 35 million reads per sample were generated. A genome index was generated, and reads were aligned to the *Lucania goodei* reference genome using STAR (version 2.7.10a) (Dobin et al., 2013). Reads mapped to exons were counted using featureCounts (version 2.0.4) (Liao et al., 2014). To further annotate genes that lacked a gene symbol after annotation of the reference genome we executed a BLASTP search using DIAMOND v2.1.20 (Buchfink et al., 2021) to the NCBI NR database (accessed December 25, 2024). When present, the topmost gene symbol was then assigned to the queried gene. These annotations were used for downstream visualization, interpretation, and gene ontology enrichment analysis.

### Statistical analysis

All analyses were conducted in R (version 4.4.0). Genes with Counts Per Million (CPM) values < 0.5 in fewer than three samples (the smallest biological replicate group size) were filtered out prior to differential expression analysis, resulting in 17,370 retained genes. Normalization was performed using the Trimmed Mean of M-values (TMM) method in edgeR (Chen et al., 2020; Evans et al., 2018). The ratio of maximum to minimum library sizes across different samples was not highly variable (∼1.5x), hence we performed differential expression analysis on log-transformed, TMM-normalized counts using the limma-trend approach in limma (Ritchie et al., 2015).

Our goals were to identify the genes involved in (a) sexual dichromatism, (b) differences between the color morphs, (c) population differences, and (d) potential genes involved in phenotypic plasticity in the expression of the blue color morph. Pairwise contrasts were computed using lmFit, constrasts.fit and eBayes functions in the limma package (Ritchie et al., 2015). First, to identify the genes involved in sexual dichromatism in both populations, we defined pairwise contrasts between males and females within each population (swamp females versus swamp males; spring females versus spring males) and examined the differentially expressed genes (DEGs) shared by both populations. To identify genes that differ across populations, we calculated contrasts between the same sex across populations (males: swamp versus spring, females: swamp versus spring) and examined the DEGs common to both sexes. We also used contrasts to identify genes involved in producing different color morphs. An initial model showed no differences between the two non-blue male morphs (red and yellow) and minimal differences between blue male morphs (blue anal fin with red pelvic and blue anal fin with yellow pelvic). Therefore, for subsequent analysis of DEGs underlying male polymorphism, we grouped the male morphs into blue and non-blue categories. We defined specified pairwise contrasts between blue and non-blue males within each population (swamp blue versus swamp non-blue, spring blue versus spring non-blue), the same morph type across populations (blue males swamp versus spring, non-blue swamp versus spring), and the interaction between morph type and population ((Blue_swamp_ vs. Non-blue_swamp_) – (Blue_spring_ vs. Non-blue_spring_)).

Next, we compared females to each of the four male color morphs within each population to examine whether all male morphs differ equally from females. We defined pairwise contrasts to identify genes that were significantly different between females and each of the four male morphs within the same population. Genes implicated in overall sexual dimorphism were consistently different between females and all four color morphs in both populations. For all comparisons, we applied a False Discovery Rate (FDR) threshold of 0.05 to identify significant DEGs.

Finally, to complement our analysis using pairwise contrasts in limma, we used a candidate-gene approach to examine whether a list of specific genes known to be involved in pigmentation synthesis, pigment cell specification, sexual dichromatism, and photoreception showed differences across our comparisons. We fit linear models to log-fold change expression data to examine the effects of sex, morph, and population on gene expression.

Significance of model terms was assessed using Type III ANOVA implemented in the Anova function of the car package, and post hoc comparisons were conducted using Tukey’s HSD (Honestly Significant Difference) via the emmeans package. The options for all analyses were set to options (contrasts = & (“contr.sum”, “contr.poly”)). Several genes were identified as significant DEGs between groups through both the limma approach with FDR correction and the linear models.

To assess the functional roles of DEGs, Gene Ontology (GO) enrichment analysis was performed using clusterProfiler (Yu et al., 2012), with gene annotations retrieved from the org.Dr.eg.db zebrafish database. GO enrichment was conducted via the enrichGO function, applying a Benjamini-Hochberg False Discovery Rate (FDR) correction threshold of 0.05 to adjust for multiple testing. We extracted GO terms for all ontologies – Biological Process, Cellular Component and Molecular Function.

## Results

### Genome Assembly

The PacBio Hifi SMRT cell yielded 1,982,991 raw reads totaling 22,567 Megabases (Mb) of sequence with a mean read length of roughly 11,380bp and an N50 read length of 12,520bp for a raw, long-read coverage of approximately 20.7x. The sequenced 10X Genomics short-read library used for ARC-based scaffolding yielded 329,944,165 pairs of reads and the sequenced Omni-C short-read library yielded 270,600,659 pairs of reads for YAHS-based scaffolding. The final assembly was 1,089,278,648bp in length, arranged in 24 chromosomes (which matches the known number of chromosomes) and 688 scaffolds with a maximum chromosome length of 49,292,181bp and an N50 scaffold length of 44,090,661bp. The BRAKER annotation yielded 22,414 protein-coding genes, arrayed in 29,482 transcripts, with BUSCO analysis reporting 95.9% completeness (6,914 complete genes, of which 217 are duplicated; Table 1).

**Table 1.**
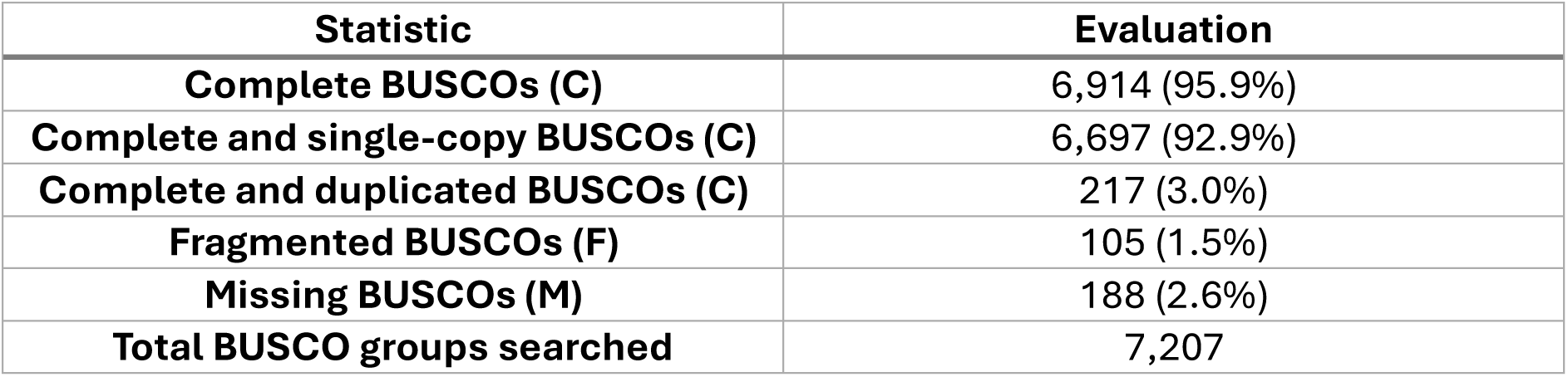
Summary of Bluefin Killifish Genome Assembly Inferred by BUSCO.

| Statistic | Evaluation |
| --- | --- |
| Complete BUSCOs (C) | 6,914 (95.9%) |
| Complete and single-copy BUSCOs (C) | 6,697 (92.9%) |
| Complete and duplicated BUSCOs (C) | 217 (3.0%) |
| Fragmented BUSCOs (F) | 105 (1.5%) |
| Missing BUSCOs (M) | 188 (2.6%) |
| Total BUSCO groups searched | 7,207 |

### Sex Differences in Gene Expression

We first sought to determine which genes underlie differences in coloration between males and females. There were 321 and 111 genes that differed between males and females from the spring and swamp populations, respectively (Figure 2a; Figure S1). Of these, 66 were common to both populations (Figure 2a; Table 2a; Table S1a). GO-term analysis indicated that these genes were enriched for pigmentation, developmental pigmentation, and melanin metabolic process (Figure 2b; Table S1b).

**Figure 2.**
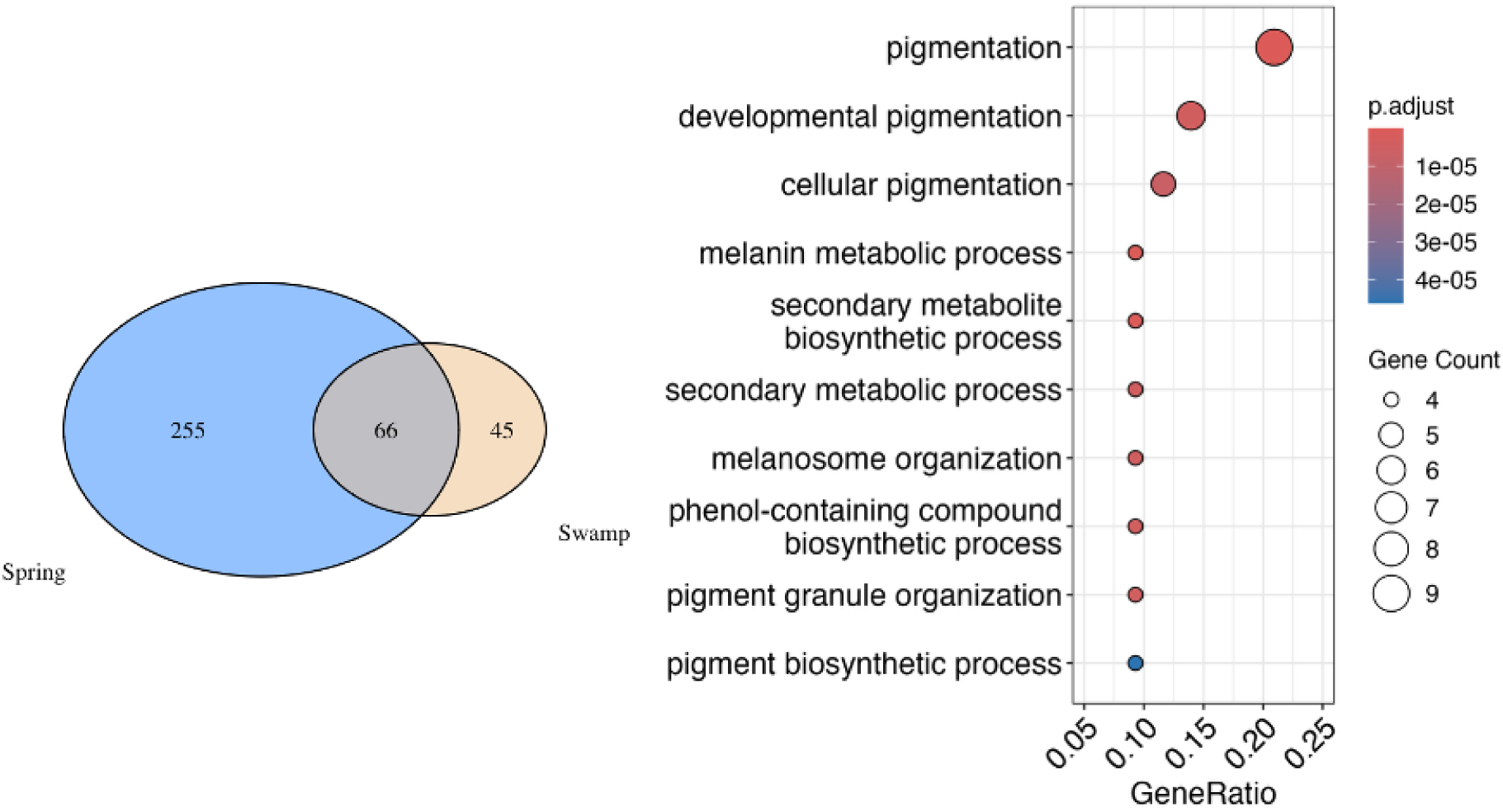
(a) The number of significantly differentially expressed genes between males and females from the two populations. Blue circle indicates the spring population, Rainbow river, and the yellow circle represents the swamp population, Everglades. (b) Results of the Gene Ontology analysis for the list of genes that significantly differ between males and females from both populations. The top 10 GO-terms for the 66 shared sexual dimorphism DEGs, across all subontology categories (biological processes, molecular function, and cellular components) are displayed here. Dot size represents the number of differentially expressed genes associated with each GO term (gene count), while dot colour indicates the adjusted P-value (Padj), with darker red colours representing more significant enrichment. The x-axis (GeneRatio) indicates the proportion of differentially expressed genes annotated to each GO term.

**Table 2.**
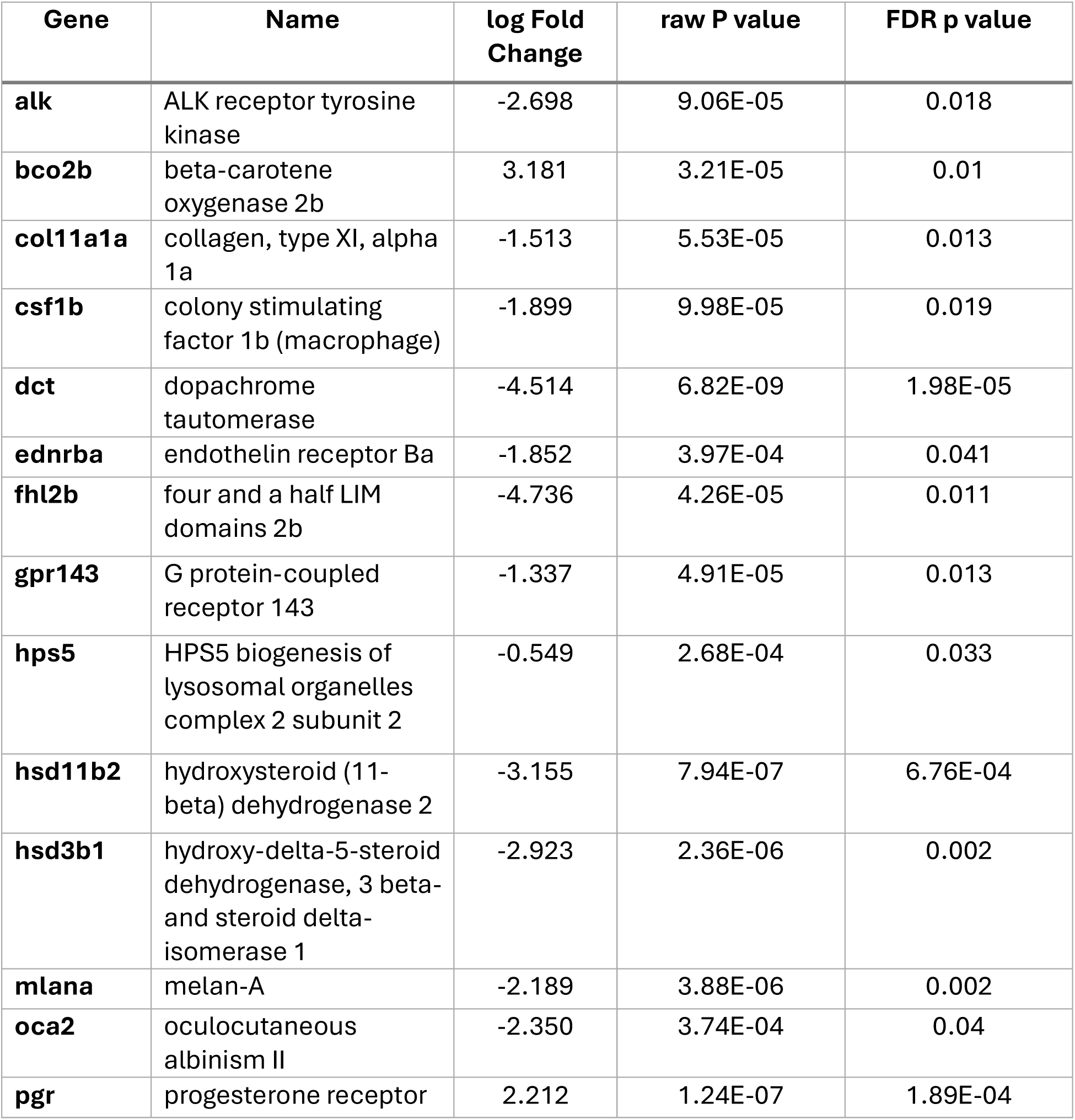
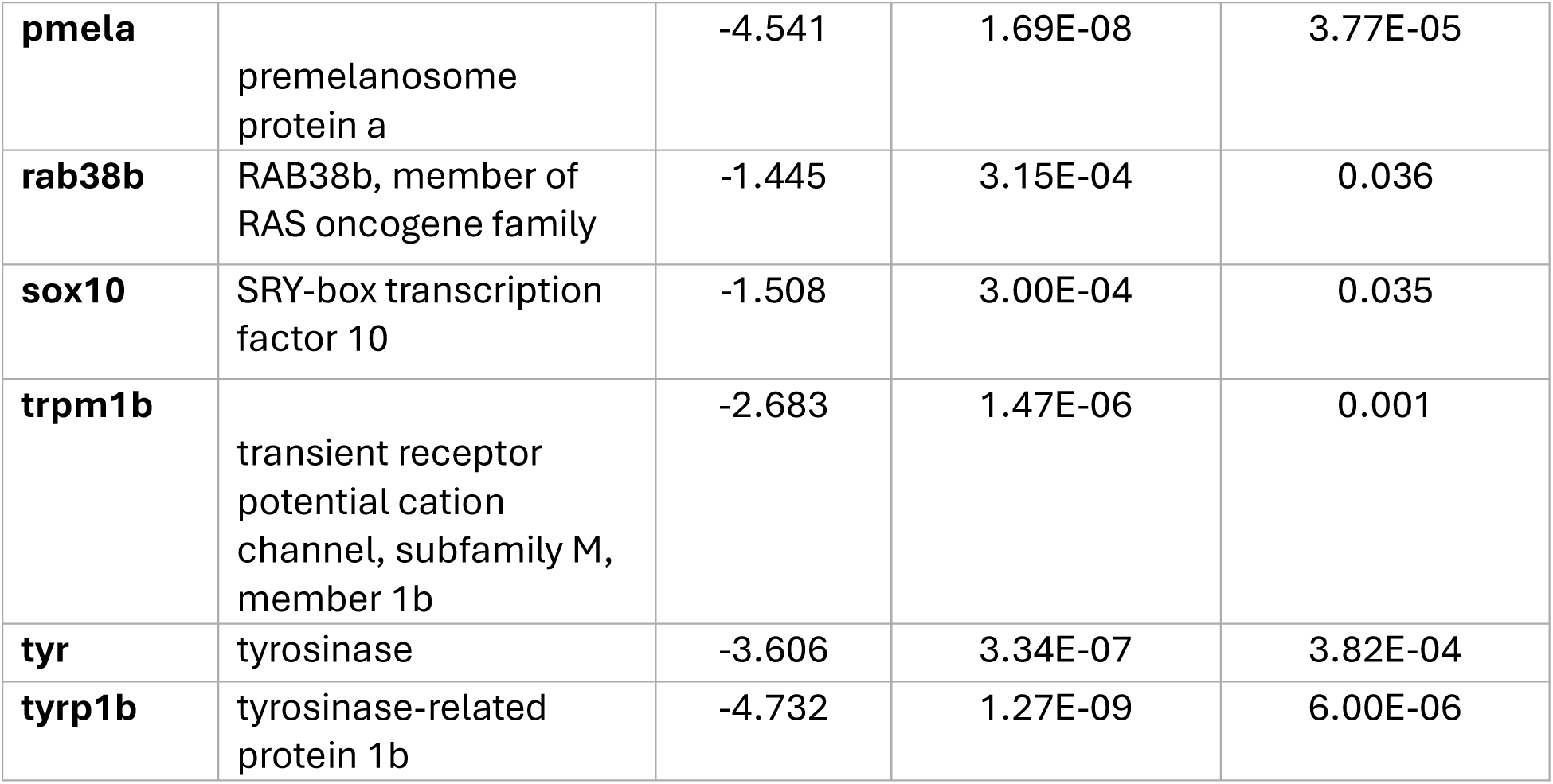
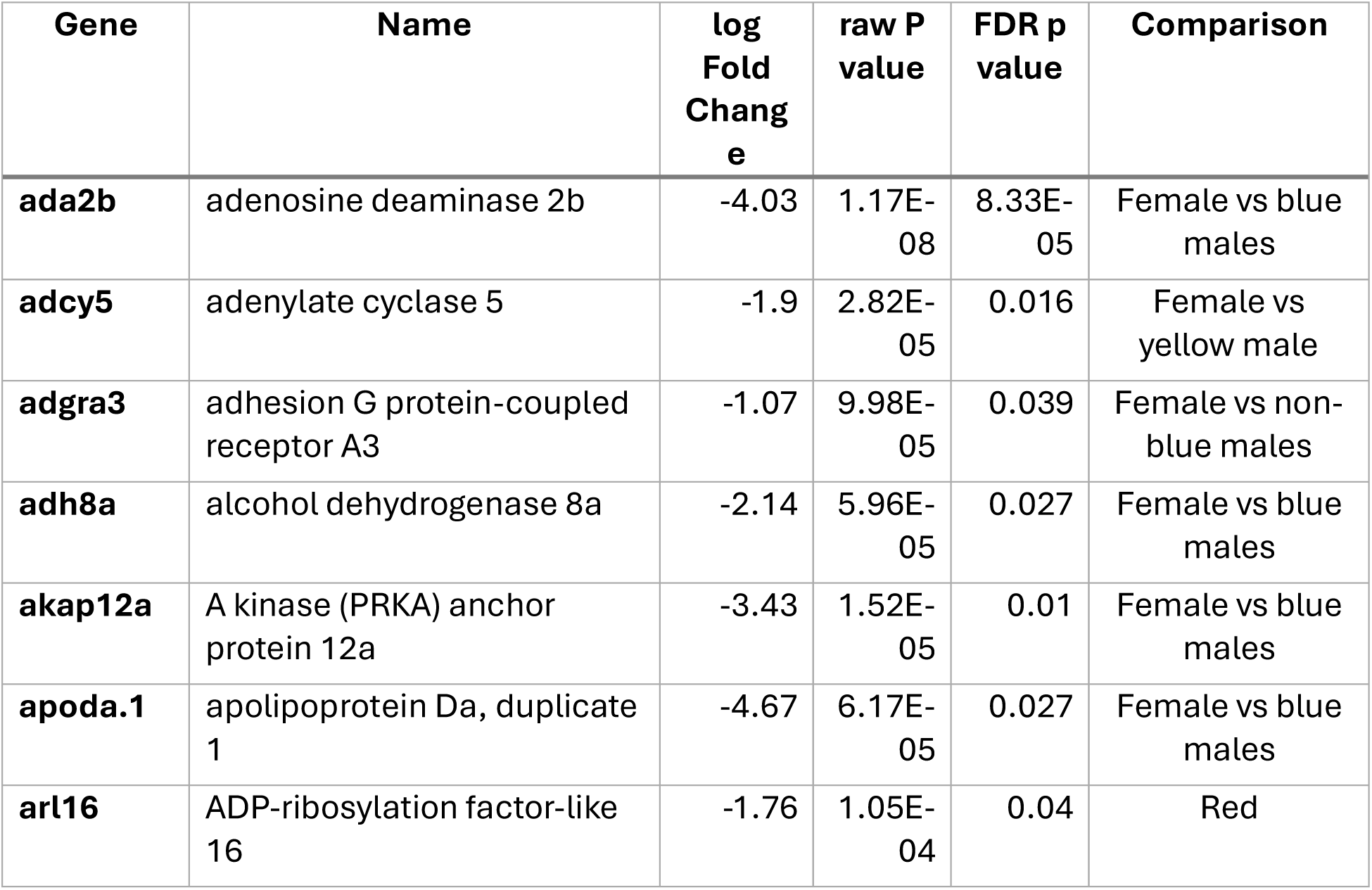
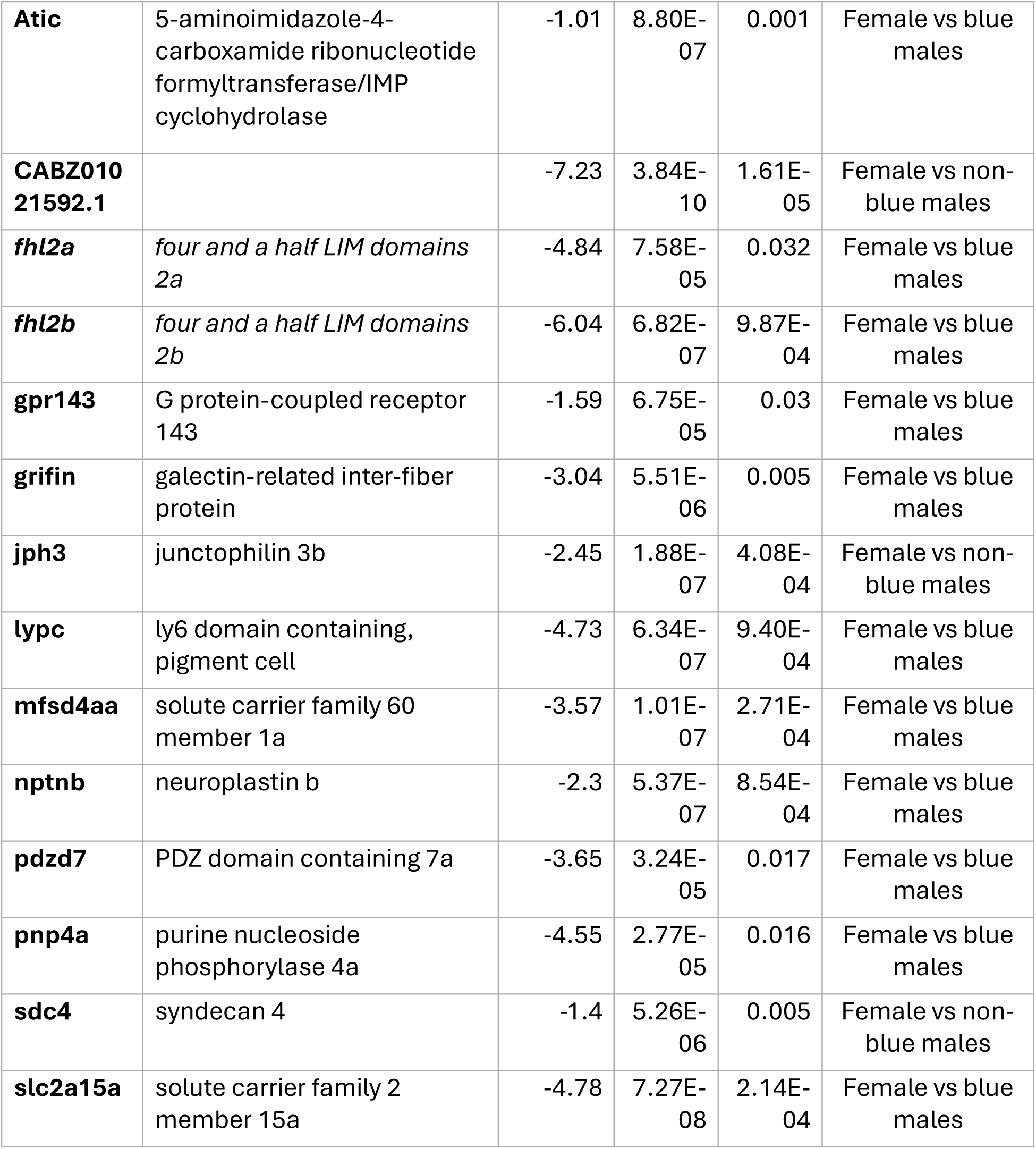
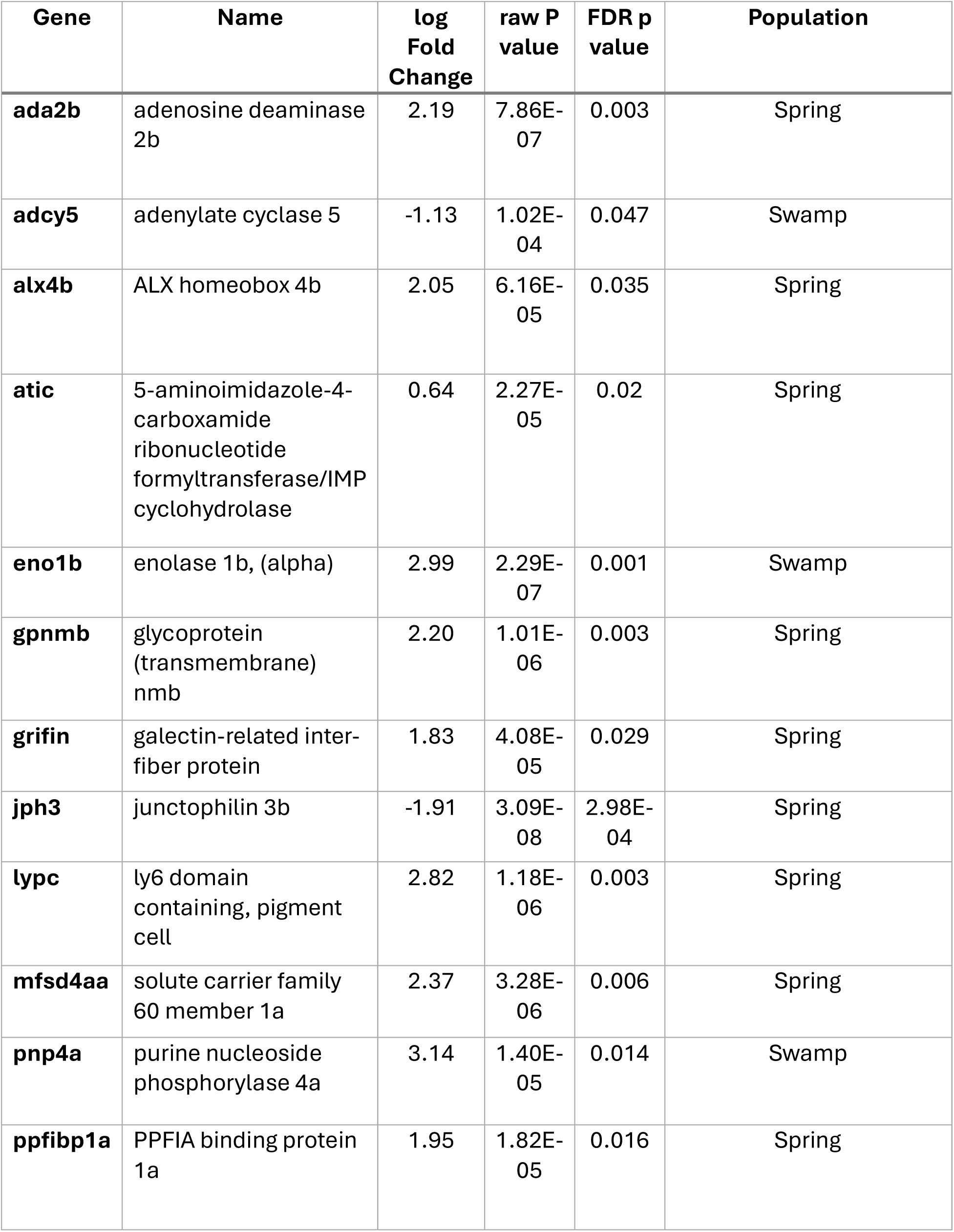
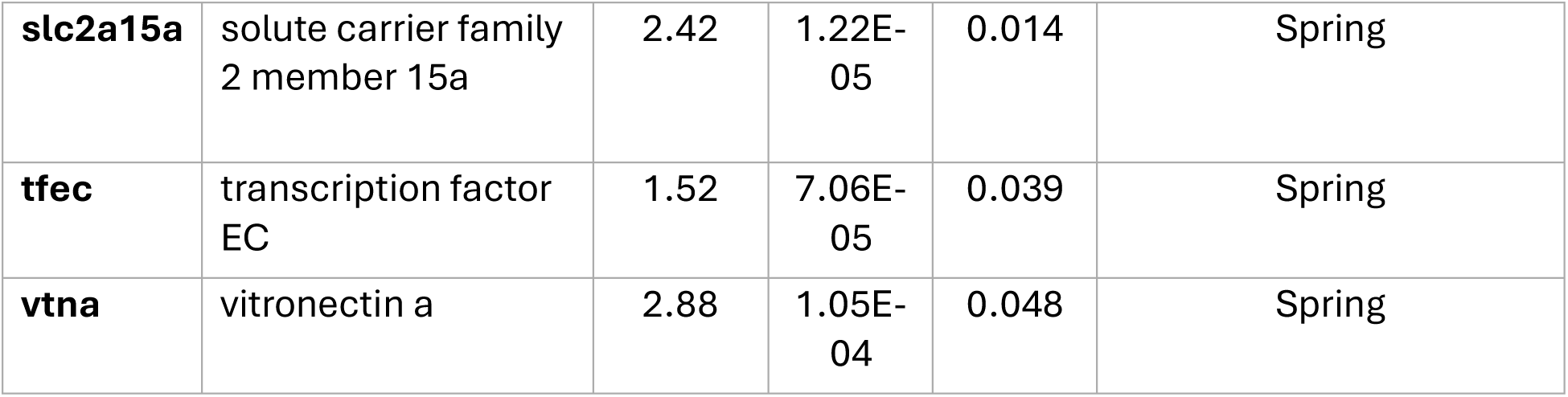
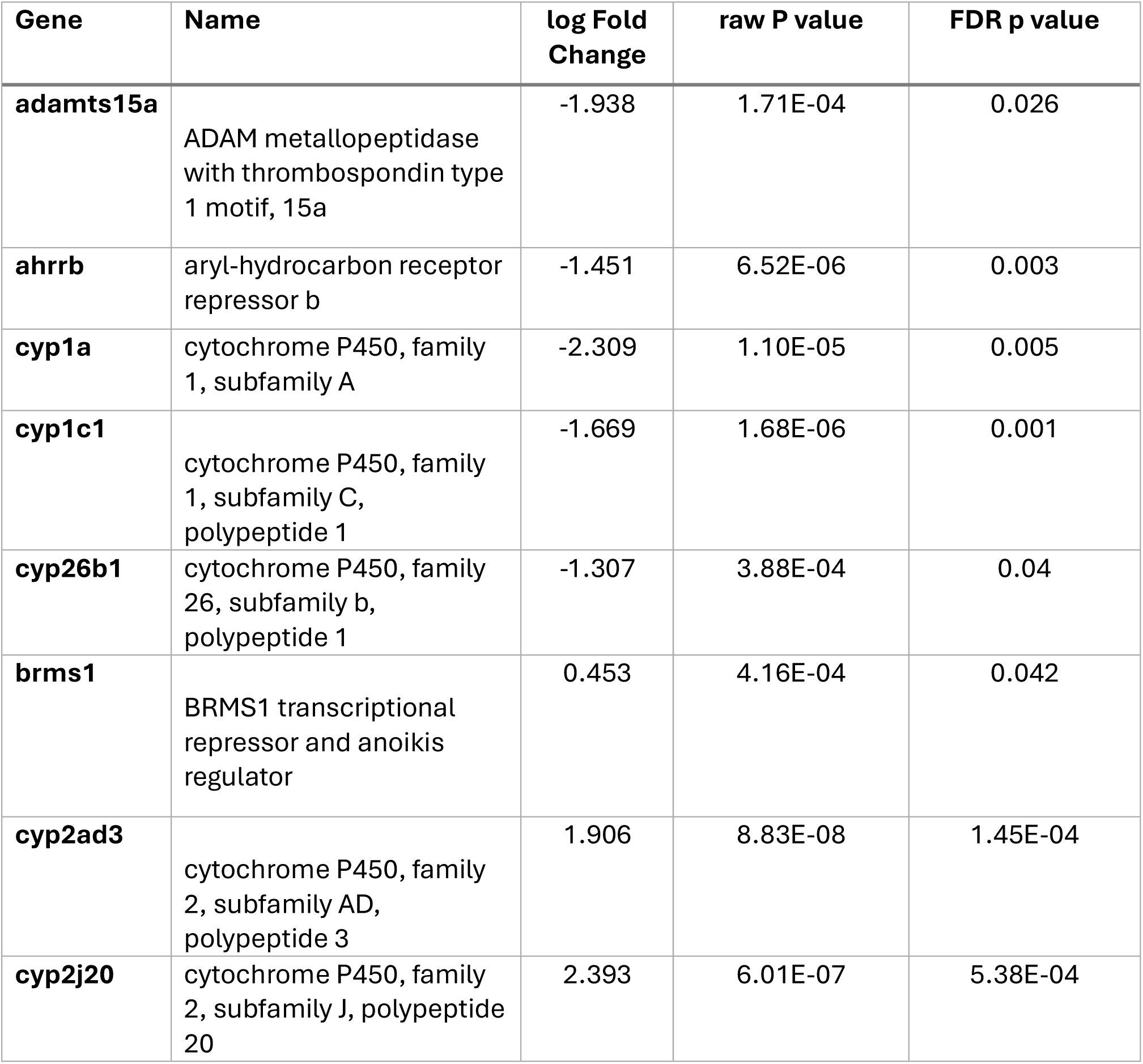
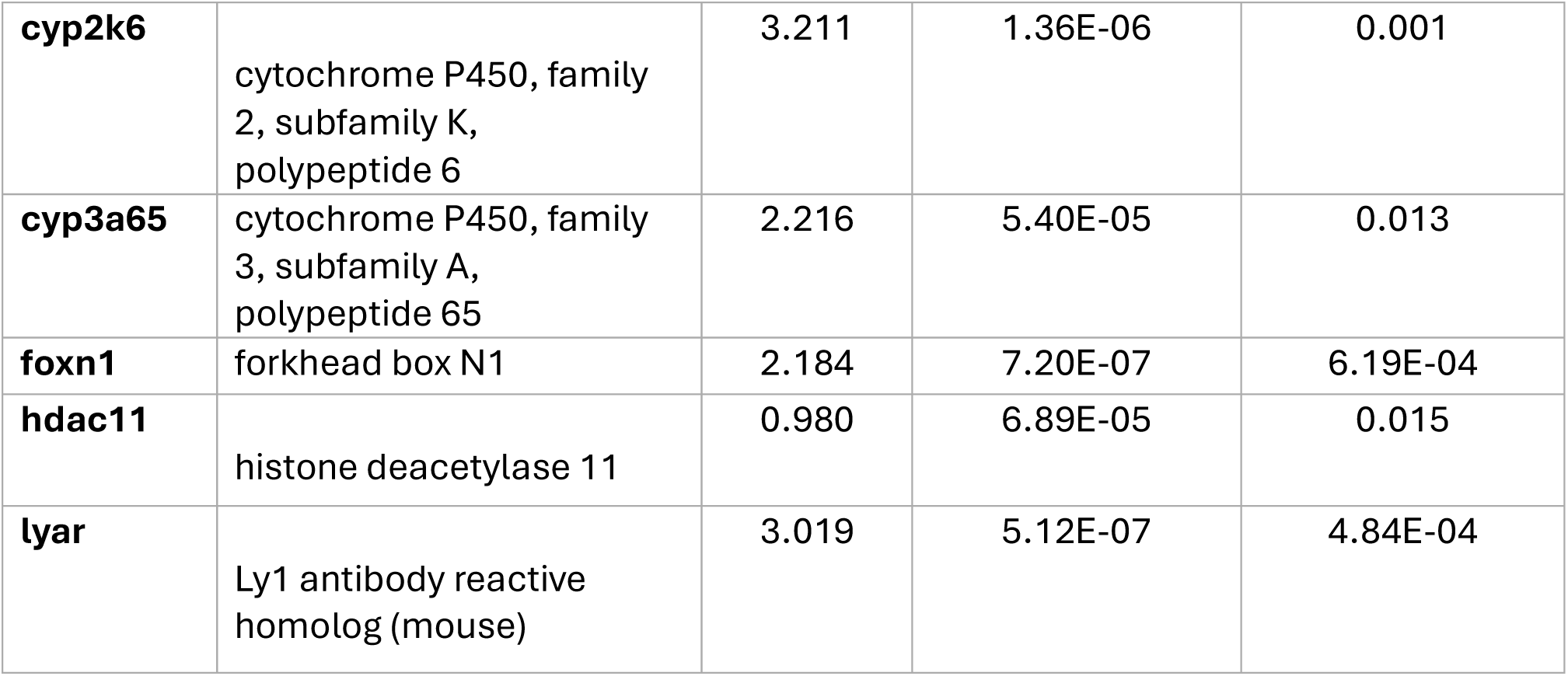
Representative genes differentially expressed in each comparison. Gene symbol, gene name, log fold change in expression, raw p value, FDR adjusted p value and description for significant DEGs from the pairwise contrast models. (a) DEGs for sex differences, (b) DEGs for each male morph vs females, (c) DEGs for blue versus non-blue males (d) DEGs for population differences between spring and swamp fish.

We also examined the number of DEGs between each male color morph and females within a population to determine whether the extent of dimorphism in gene expression was uniform across all morphs (Table S2a; Table S2b). Figure 3 displays DEGs differentially expressed between each color morph and females for both populations. We found that all four morphs shared a set of DEGs that differed from females (Figure 3; Table 2b). Among the shared DEGs from both analyses, we found that many canonical genes involved in melanin synthesis were upregulated in males compared to females, including enzymatic genes (*tyr, dct, oca2, pmela, tyrp1*) and a solute carrier (*slc45a2*) (Figure 4).

**Figure 3.**
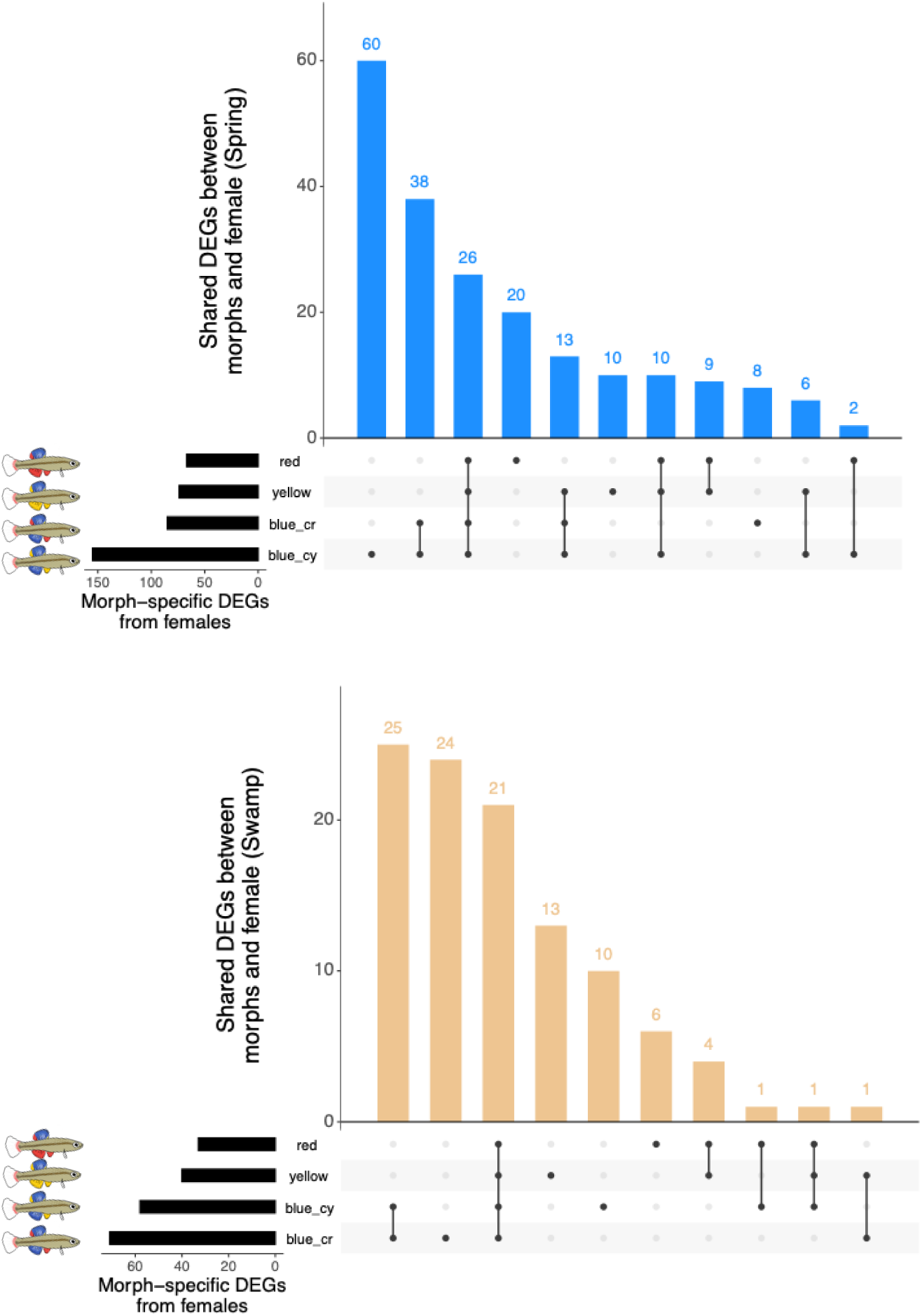
UpSet plots showing the overlap in differentially expressed genes (DEGs) between male morphs compared to females in (a) spring and (b) swamp populations. Horizontal bars (left) indicate the total number of DEGs identified for each male morph. The connected dot matrix (bottom) indicates the morph(s) represented in each intersection, with filled and connected dots denoting shared gene sets and single filled dots denoting morph-specific gene sets. Vertical bars (top) indicate the number of DEGs within each unique or shared intersection. Blue_cy - blue male morphs with yellow pelvic fins; blue_cr – blue male morphs with red pelvic fins.

**Figure 4.**
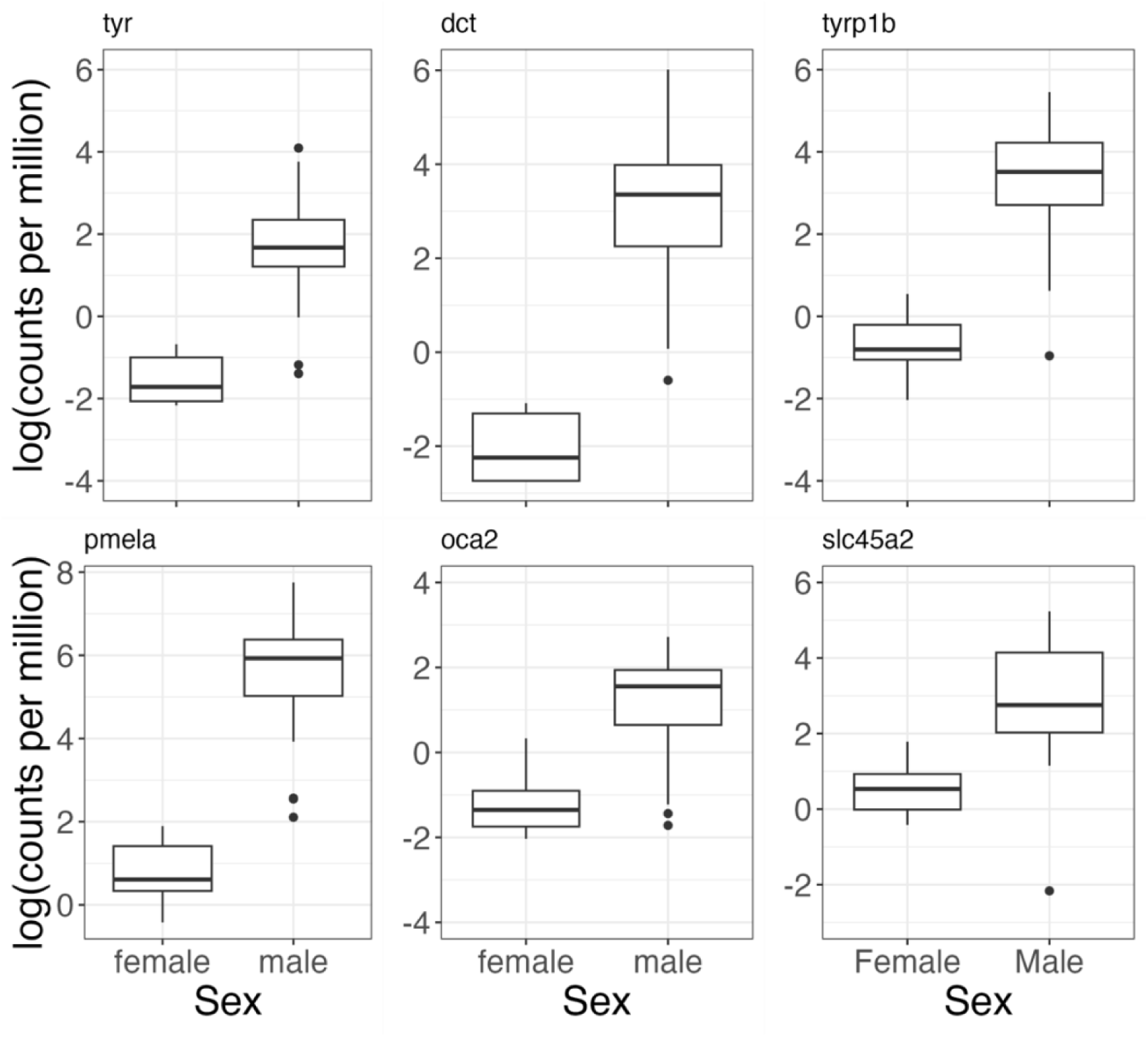
Pigmentation genes that differ significantly as a function of sex. Log counts per million expression of the genes coding for the enzymes tyrosinase (*tyr*), DOPAchrome tautomerase (*dct*), Tyrosinase-related protein 1b (*tyrp1b*), Premelanosome protein a (*pmela*), transmembrane protein oculocutaneous albinism II (*oca2*), and solute carrier *slc45a2*. These genes code for proteins that are crucial in the melanin biosynthesis pathway.

We used candidate gene analysis to determine whether any of the common sex steroid receptors differed between males and females (Table 3). The progesterone receptor gene, *pgr,* was the only significant DEG between males and females and was significantly upregulated in females (Figure 5). We found no evidence of differential expression of androgen or estrogen receptors between the sexes (Figure 5).

**Figure 5.**
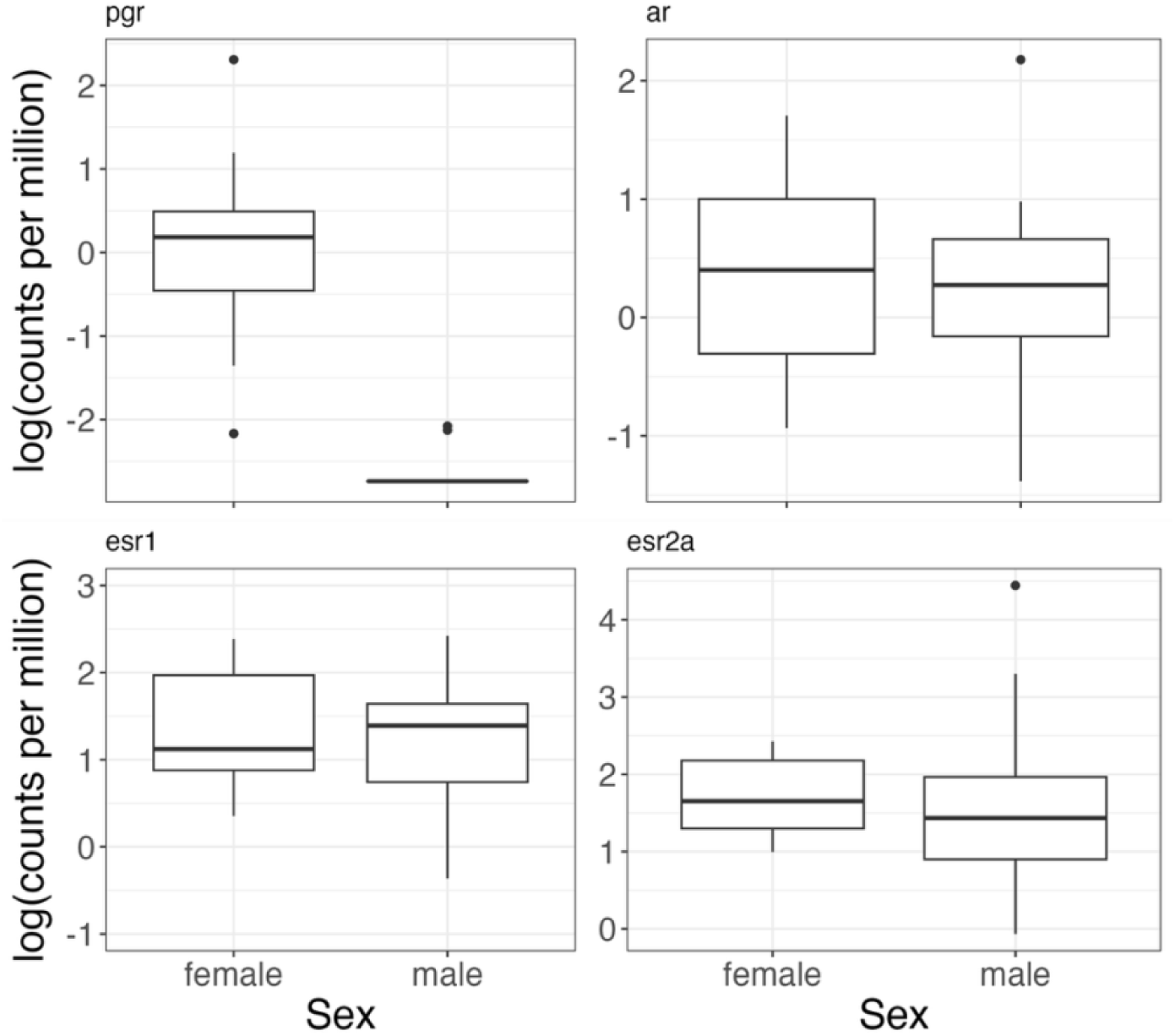
Sex steroid receptor expression as a function of sex. Box plots show log counts per million expression of the genes coding for the following sex steroid hormone receptor proteins – progesterone receptor (*pgr*), androgen receptor (*ar*), estrogen receptors *esr1* and *esr2a*. Significant sex difference in expression is seen only in the progesterone receptor gene, *pgr*.

**Table 3.**
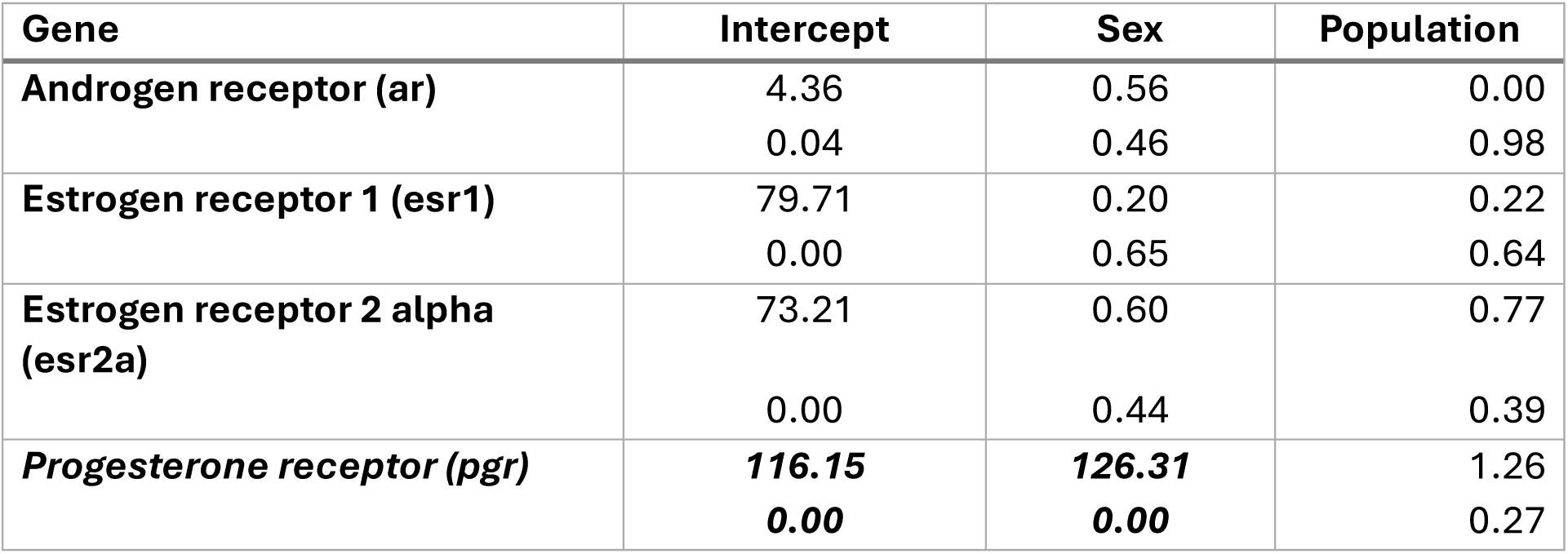
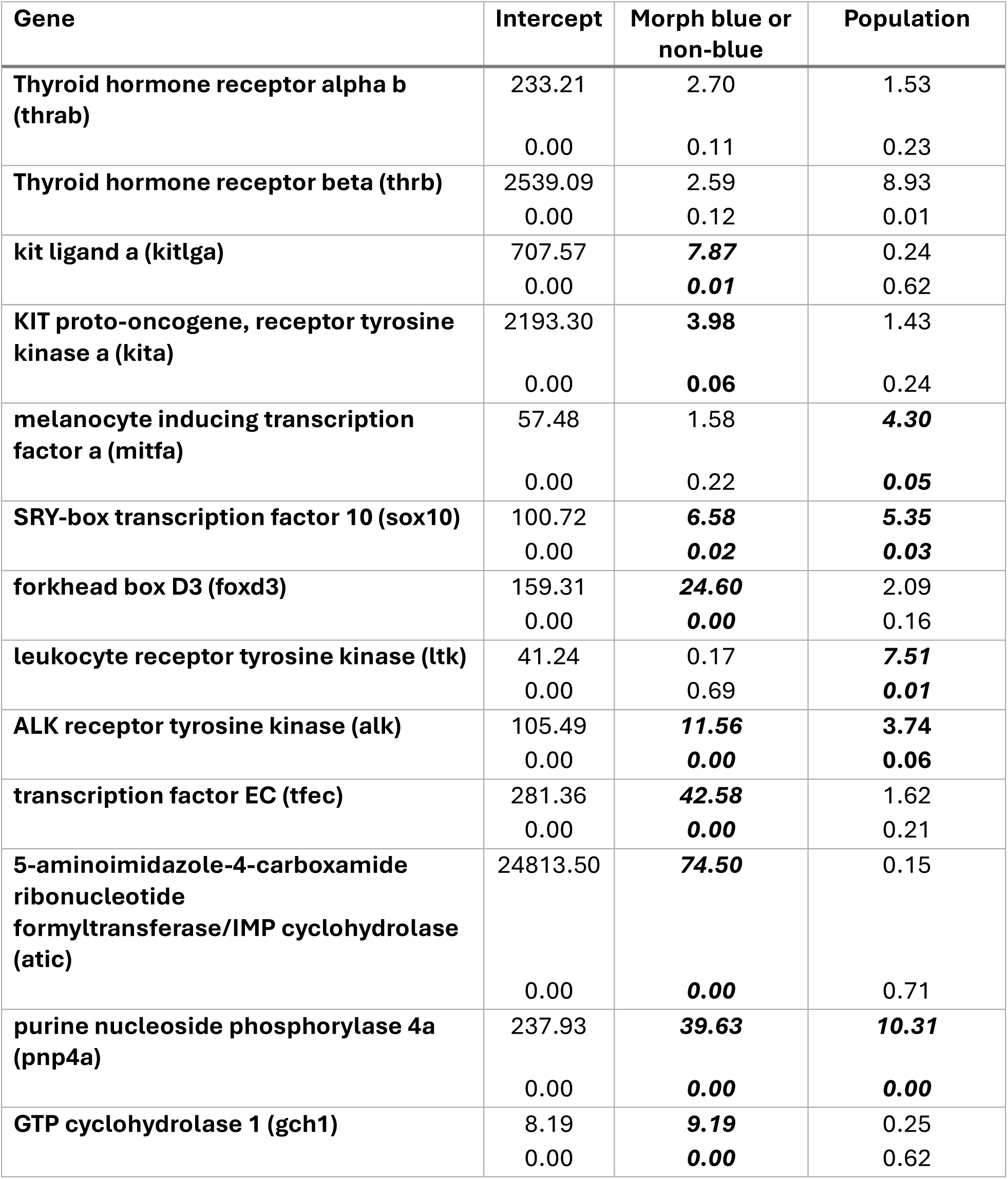
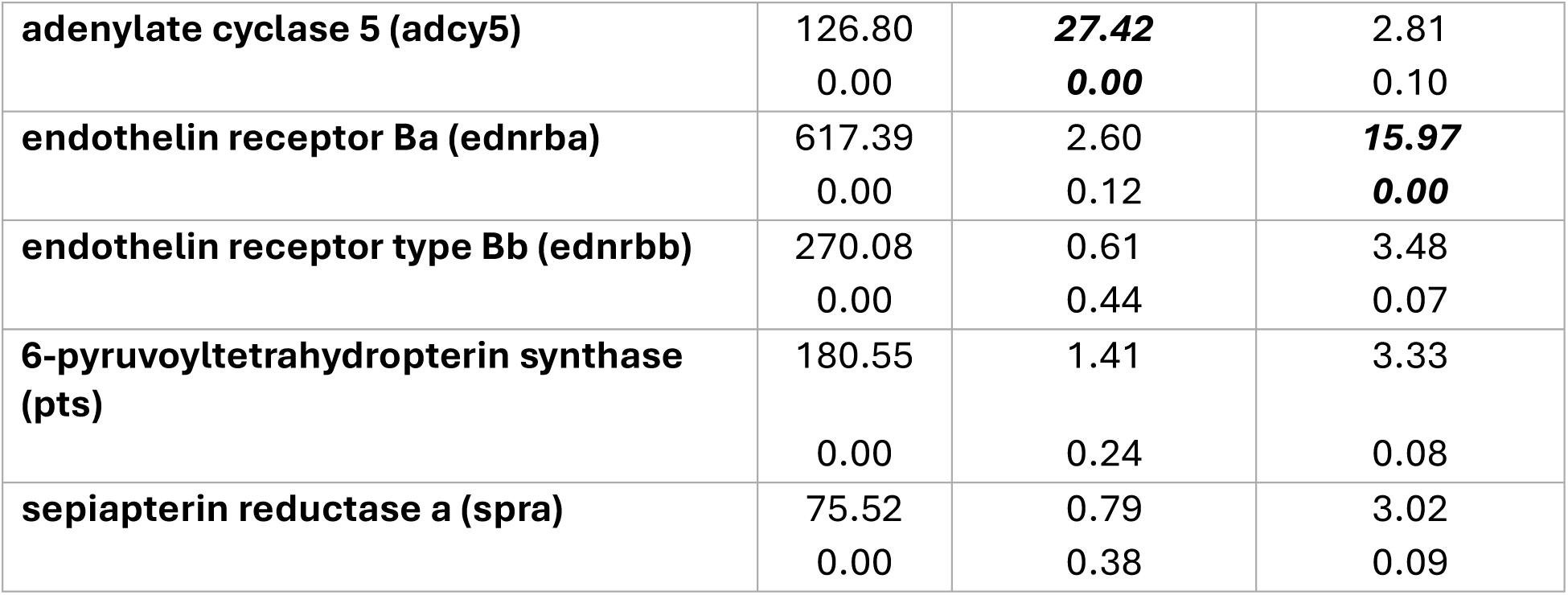
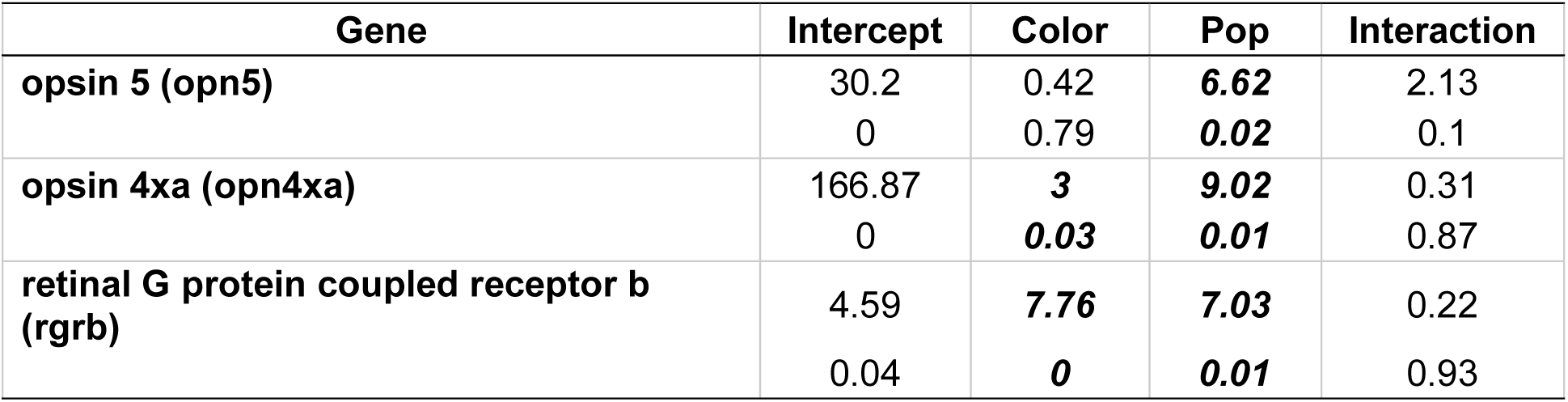
Type 3 analysis of variance on the log-fold change in gene expression of sex steroid hormone receptor genes (a), male pigmentation genes (b), non-visual opsin genes (c). a. Analyses of variance as a function of sex and population for gene expression in log counts per million. F-values are listed on top and p-values below. Values in bold and italic show statistically significant effects of sex on gene expression. Denominator DF are 36 for all models. **Table 3. b.** Analyses of variance as a function of male blue morph status and population for gene expression in log counts per million . F-values are listed on top and p-values below. Values in bold and italic show statistically significant main effects of male morph and population on gene expression. Denominator DF are 28 for all models. **Table 3. c.** Analyses of variance as a function of fin color (sex and male morph), population and their interaction for gene expression in log counts per million. F-values are listed on top and p-values below. Values in bold and italic show statistically significant main effects of fin color, population and their interaction on gene expression. Denominator DF are 29 for all models.

### Color Morph Differences in Gene Expression

We next compared the number of DEGs between each color morph and females of each population to assess patterns in the total number of differentially expressed genes across color morphs. Blue male morphs had a larger number of DEGs that differed from females than non-blue males (Figure 3, horizontal bars) as well as a higher number of unique DEGs from females (Figure 3, vertical bars). Both populations showed similar patterns of sexual dimorphism in gene expression with respect to blue and non-blue male DEGs, although spring population fish showed a higher number of DEGs for each comparison (Table S2a; Table S2b). We found no DEGs between red and yellow males in either population after FDR correction. Likewise, we found few DEGs between blue males with yellow pelvic fins versus blue males with red pelvic fins. We therefore condensed the analysis to compare blue versus non-blue males.

Both blue morph status and source population had strong effects on gene expression, but there was no interaction between them (Figure S2; Table S3a). There were 38 and 31 genes that differed between blue and non-blue males from the swamp and spring populations, respectively, of which 20 genes were shared between the populations (Figure S3). The shared DEGs included genes involved in the iridophore synthesis and differentiation pathways (*atic,* and *pnp4a*; Figure 6, Table S3b), which were significantly upregulated in blue male morphs compared to non-blue males. Additionally, a transcription factor influencing iridophore specification, *tfec*, was significantly upregulated in blue male morphs, indicating a putative candidate gene involved in the plastic blue anal fin coloration (Figure 6).

**Figure 6.**
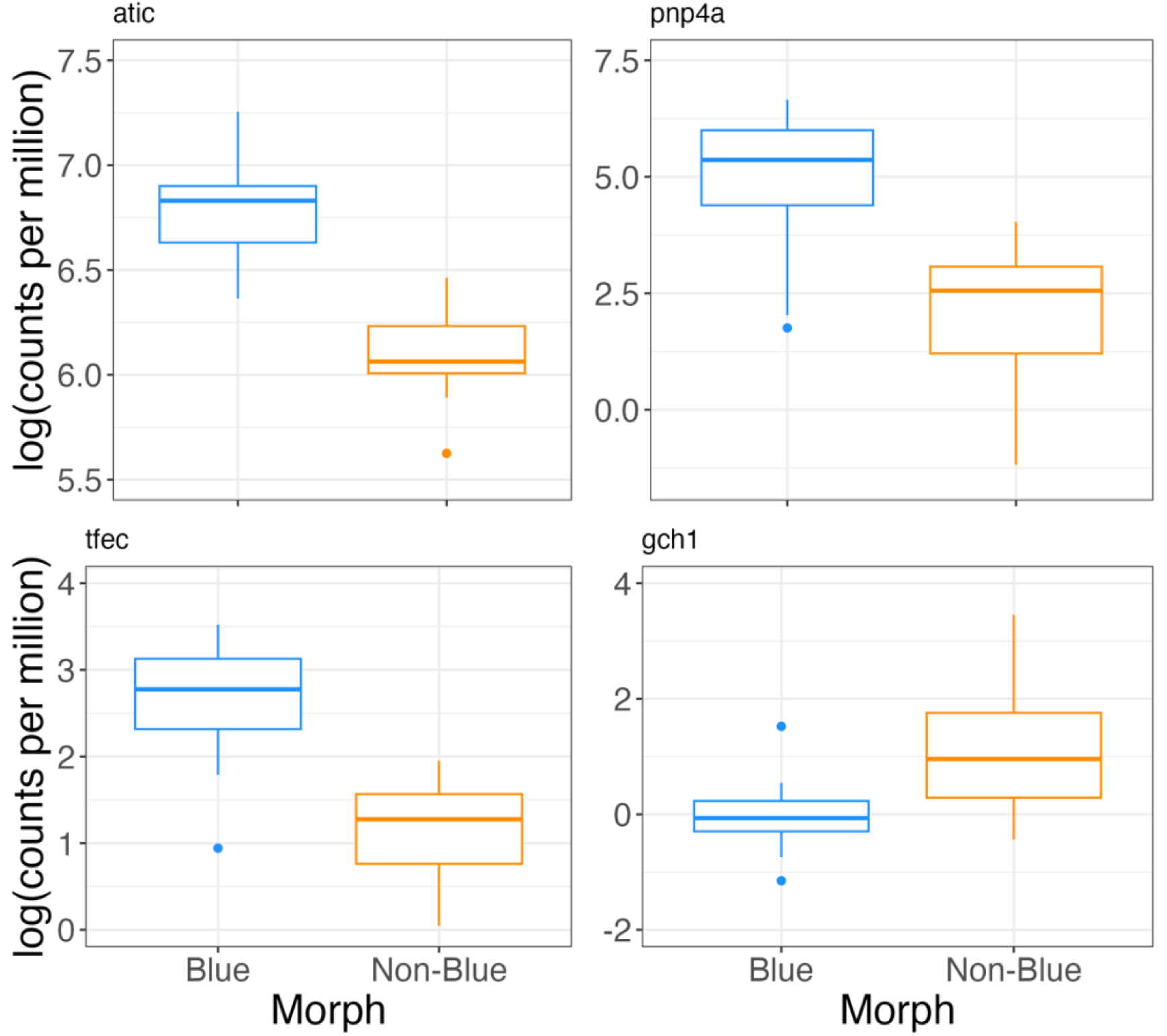
Gene expression differences between blue and non-blue male morphs. Bpx plots show log counts per million expression of the genes *atic* (5-aminoimidazole-4-carboxamide ribonucleotide formyltransferase), *pnp4a* (purine nucleoside phosphorylase 4a), *gch1* (GTP cyclohydrolase 1) and *tfec* (transcription factor EC). Values for males of both populations are shown here.

Based on work in zebrafish (Irion & Nüsslein-Volhard, 2019; Patterson & Parichy, 2019), we hypothesized that many of the genes involved in the developmental switch between iridophores and xanthophores (as well as iridophore specification) would be differentially expressed between blue and non-blue males. Through candidate gene analysis, we examined the influence of male morph status on thyroid hormone receptors, chromatophore transcription factors, and pteridine pigment pathway (producing red and yellow pterins) enzymes previously implicated in the regulation of chromatophores (Hashimoto et al., 2021; Kelsh et al., 1996; Parichy, 2006b; Saunders et al., 2019). In addition to the transcription factor *tfec* identified through pairwise contrasts, candidate gene analysis revealed a significant effect of morph status on several transcription factors implicated in iridophore specification and cell fate determination (*kitlga, kita, sox10, foxd3, alk*; Table 3). We did not find a significant effect of male morph status on thyroid hormone receptor genes (*thrab, thrb;* Table 3), although there was a strong effect of population, where spring fish had higher expression levels of *thrb* than swamp fish. This pattern is in keeping with the role of thyroid hormone in suppressing iridophore development. The population pattern reflects the difference in means between non-blue and blue fish, with non-blue fish having higher expression levels of both *thrb* and *thrab* than blue fish, although the differences were not statistically significant (*p∼*0.12 for both tests).

Far fewer genes (seven) were upregulated in non-blue males. We found that the *gch1* gene (GTP cyclohydrolase 1, Figure 6), which is involved in the first step of the pteridine synthesis pathway, was significantly upregulated in non-blue (i.e., red and yellow) males (Braasch et al., 2007b; Hashimoto et al., 2021; Ziegler, 2003). However, candidate gene analysis found no evidence for upregulation of any other genes involved in pterin synthesis. While many of the canonical genes for pterin synthesis were present, they were not differentially expressed.

### Overall Population Differences

Several genes differed significantly as a function of population irrespective of sex (Table S4a). GO term enrichment analysis indicated that these genes are broadly categorized under the molecular function domain, with specific GO terms indicating enzymatic and ion-binding functions (Table S4b). Although the specific roles of these genes in lighting environment or population differences are not well-studied, we found that several genes encoding enzymes from the cytochrome P450 family as well as certain histone deacytelation genes were upregulated in swamp population fish (Table 2d).

### Light-Sensitive Candidate Genes

Lastly, we used the candidate gene approach to examine the expression of extraocular opsin genes. We found significant main effects of fin color and population on retinal G protein-coupled receptor b (*rgrb*) and opsin 4xa (*opn4xa*), but no effect of their interaction (Table 3). Blue males from both populations showed significantly higher levels of retinal G protein-coupled receptor b (*rgrb*) compared to females and non-blue males (Figure 7), and males showed significantly higher levels of expression of melanopsin, *opn4xa* (Figure 7). We also saw a significant main effect of population on the expression of opsin 5, with swamp fish showing higher expression of this gene (*opn5*; Table 3).

**Figure 7.**
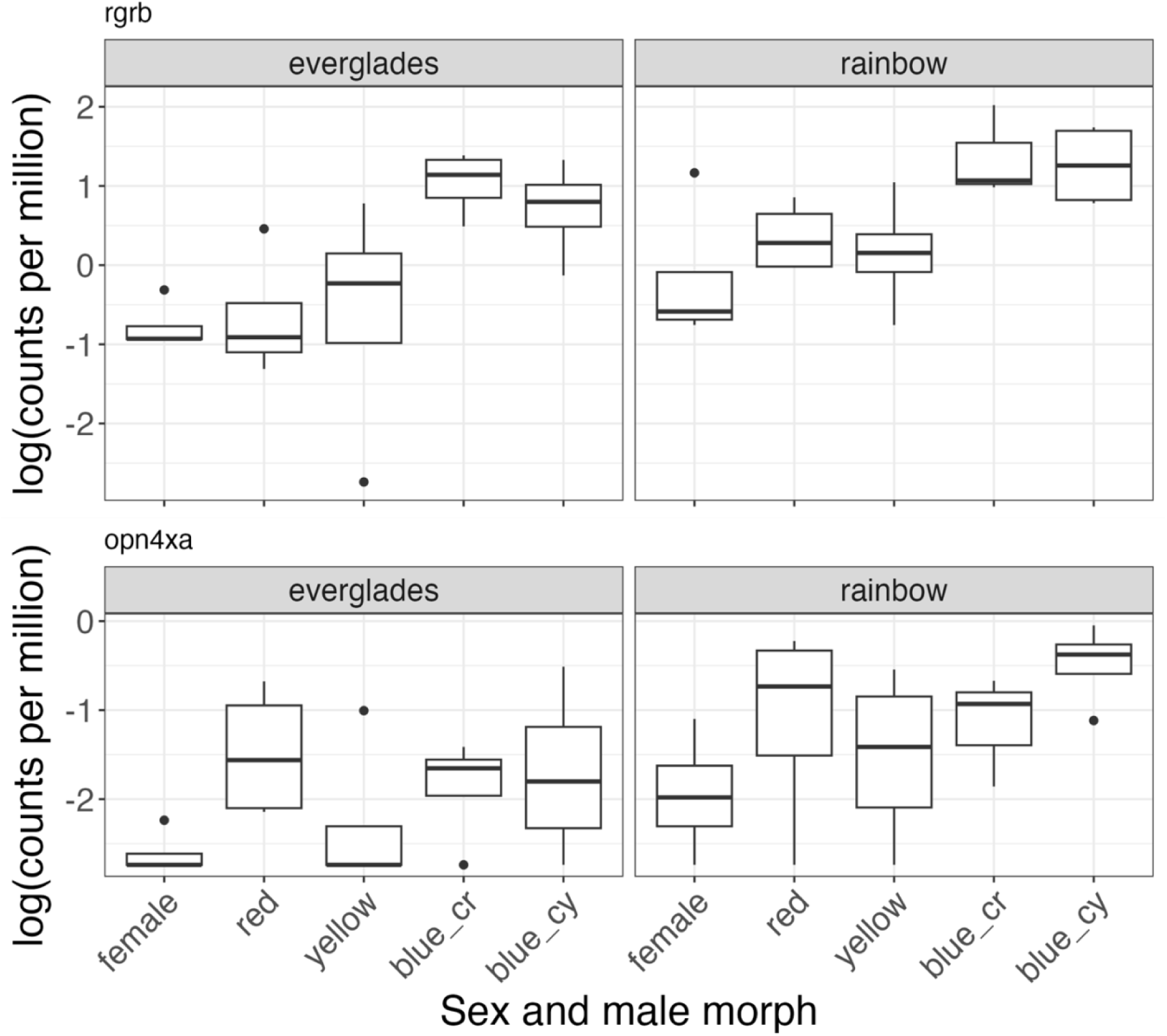
Expression of extraocular, non-visual opsin genes. Box plots show log counts per million expression of the non-visual opsin genes (top) *rgrb* (retinal pigment epithelium (RPE) retinal G protein receptor 2) and (bottom) *opn4xa* (opsin 4xa, a melanopsin) as a function of male color pattern and population.

## Discussion

Bluefin killifish harbor extensive variation in coloration at multiple levels: between the sexes, between blue versus non-blue males, and between yellow and red pterin-producing males (Figure 1). In this study, we examined the transcriptomic differences underlying sexual dichromatism and male color polymorphism in anal fin tissue. We found striking variation between males and females as well as between blue and non-blue males. Many of the differentially expressed genes are known to play important roles in color production, specifically melanin and iridophore production.

Gene expression varied as a function of sex, and many of these genes involved the upregulation of melanin synthesis genes in males compared to females. Every step of the melanin biosynthesis pathway (*tyr, dct, tryp1, pmela*), as well as membrane transport proteins responsible for transporting the substrates of eumelanin (*oca2, slc45a2*) (Braasch et al., 2007a; del Marmol & Beermann, 1996; Hashimoto et al., 2021; Kelsh et al., 1996), were upregulated in males. These sex differences in melanin pigmentation gene expression are consistent with our understanding of the melanin biosynthesis pathway, as inferred from studies using mutant and knockout zebrafish and medaka (Dooley et al., 2013; Hashimoto et al., 2021; Wang et al., 2022). Our analysis most likely had high power to detect these differences because all males have a black border around the anal fin. In bluefin killifish, variation in melanin on the anal fin is thought to function as a signal of social dominance status (Johnson & Fuller, 2015). The consistent upregulation of melanin-related genes in all males relative to females may reflect the dynamic nature of this trait, suggesting that melanin synthesis remains active in adult males.

Hormonal regulation often underlies sex-limited trait expression (Williams & Carroll, 2009; Mank, 2023), and androgens play a role in the expression of male coloration in bluefin killifish (Fuller & Travis 2004). In many fish species, males and females differ not only in circulating androgen levels but also in androgen receptor expression in peripheral tissues involved in color, morphology, and behavior (Carver et al., 2021; Forlano et al., 2010; Ogino et al., 2023; Ryu et al., 2024; Schuppe et al., 2017). We expected to find differences in the expression of androgen receptor genes or in the expression of enzymes involved in androgen synthesis. In contrast, we found that the progesterone receptor was the only sex steroid receptor differentially expressed between the sexes, despite the presence of other sex steroid receptors in the anal fins of males and females (Figure 5). One possibility is that males have higher levels of circulating androgens (Angus et al., 2001; Hopper, 1949; Sangster, 1948; Turner, 1942), driving sexual dimorphism even though androgen receptors themselves do not appear to differ in abundance. Consistent with this hypothesis, androgens induce male coloration in females, and juvenile males initially resemble females in coloration (Foster, 1967; Fuller & Travis, 2004). Additionally, elevated progesterone receptor expression in females might be antagonistic to androgen-responsive pathways, resulting in the downregulation of male coloration genes and the maintenance of clear fins. Together, these findings highlight promising directions for future research on the hormonal regulation of dichromatism in bluefin killifish.

In addition to the shared set of differentially expressed genes between males and females, there were also differences in the number of DEGs between each male color morph and females. Blue males exhibited many more sexually dimorphic DEGs and a substantially larger set of morph-specific expression differences compared to non-blue males, indicating additional layers of regulatory and phenotypic divergence. Blue and non-blue male coloration differ fundamentally in their mechanistic basis, with blue coloration arising from structural mechanisms involving the precise arrangement of reflective platelets produced as a byproduct of the guanine synthesis pathway (Braasch et al., 2007b; Hashimoto et al., 2021; Irion & Nüsslein-Volhard, 2019). The increased number and uniqueness of DEGs in blue males may therefore reflect the greater regulatory complexity required to generate and maintain structurally based coloration, especially since it is thought to be more environmentally responsive (Kemp & Rutowski, 2007; McGraw et al., 2002).

Among the few genes that were upregulated in non-blue males, we found that a gene encoding a key, rate-limiting enzyme upstream of the pteridine synthesis pathway, *gch1* (GTP cyclohydrolase 1) (Ziegler, 2003), was significantly upregulated (Figure 6). Surprisingly, genes encoding enzymes acting downstream in the pteridine synthesis pathway (Table 3b)—*pts* (6-pyruvoyltetrahydropterin synthase) and *spra* (sepiapterin reductase a)—did not differ significantly between blue and non-blue males. Furthermore, we found no differences in gene expression between red and yellow males. Red males produce both drosopterin and xanthopterin, whereas yellow males express only xanthopterin (Johnson & Fuller, 2015).

Hence, our expectation was that there would be differences in genes related to drosopterin synthesis, which we did not find. The inheritance of red versus yellow coloration suggests that an autosomal locus of large effect is segregating in all known bluefin killifish populations.

Taken together, the results suggest that variation in red versus yellow anal fin coloration does not involve large-scale disruption of the pteridine synthesis pathway.

A larger number of genes were upregulated in blue morphs, suggesting that the production of the blue coloration involves more steps. Blue males showed significant upregulation of genes involved in purine metabolism and production of guanine reflective platelets (*atic*, 5-aminoimidazole-4-carboxamide ribonucleotide formyltransferase and *pnp4a,* purine nucleoside phosphorylase 4a), and a number of transcription factors involved in iridophore specification (*tfec* - transcription factor EC; *kitlga* - kit ligand a; *kita* - KIT proto-oncogene, receptor tyrosine kinase a; *sox10* - SRY-box transcription factor 10; *foxd3* - forkhead box D3). Research using mutant zebrafish has established that these transcription factors not only influence downstream iridophore specification genes, but also influence each other through feedback loops (Curran et al., 2010; Higdon et al., 2013; Kelsh, 2004; Parichy, 2006b; Patterson et al., 2013; Wang et al., 2023). Interestingly, the transcription factor EC (*tfec*) has been proposed to play a particularly pivotal role in directing iridophore specification from multipotent neural crest progenitors during development (Petratou et al., 2018, 2021) and is thought to be a master regulator of iridophore fate. In this study, *tfec* was the transcription factor with the most robust pattern of differential expression between blue and non-blue male morphs, indicating that it could potentially be a key component of the plasticity mechanism underlying the blue phenotype.

One of our a priori predictions was that thyroid hormone signaling might be important to the creation of blue versus non-blue male morphs. Thyroid hormone is known to influence the relative amounts of black and orange coloration in guppies and other fish (Liu et al., 2024; Prazdnikov, 2021, 2025). Thyroid hormone receptor protein has also been shown to influence cell specification of chromatophores and iridophores and can have an inhibiting effect on iridophore development (McMenamin et al., 2014; Saunders et al., 2019). Our results were mixed. There was no statistically significant effect of blue-male status on thyroid receptor levels, although, on average, non-blue males had higher expression levels of the thyroid receptors *thr* and *thrab* than blue males. Thyroid hormone receptor *thrb* was significantly higher in the spring population, which has fewer blue males on average. In addition, we did find a significant effect of male morph status on several downstream targets of thyroid hormone and thyroid receptor proteins. These include genes involved in melanin (colony stimulating factor 1r, tyrosinase, DOPAchrome tautomerase, solute carrier 24a5), iridophore (ALK receptor tyrosine kinase (*alk*), leukocyte receptor tyrosine kinase (*ltk*)) and pterin (GTP cyclohydrolase 1 (*gch1*)) pathways. Receptor-level differences may simply be more transient or restricted to specific developmental windows than the broader transcriptional signature we detected downstream.

Another potential mechanism for the creation of blue versus non-blue males is that they differ in the expression of non-visual opsin in the fins themselves. Past work has shown that males are more likely to express blue coloration when raised in swamp-mimicking, tannin-stained environments, which suggests that detection of the lighting environment is likely important.

The role of extraocular, non-visual opsins has recently gained attention as a potential mechanism mediating environmentally induced changes in coloration (Cronin & Johnsen, 2016; Davies et al., 2015; Policarpo et al., 2025; Schweikert et al., 2023). We found that blue male morphs significantly upregulate *rgrb* (retinal pigment epithelium [RPE] retinal G protein receptor 2). Little is known about *rgrb* as a nonvisual opsin. One study has shown that it was weakly expressed in the fin tissue of a flounder (Liu et al., 2020). To the best of our knowledge, no previous study has found differential expression of this nonvisual opsin as a function of sex or color morph. Similarly, males also have significantly higher expression of opsin 4xa in comparison to females. Opsin 4xa is a well-known extraocular opsin important for not only photoreception, but also light-sensitive physiological processes (Davies et al., 2015; Provencio et al., 1998). Sex differences in opsin 4xa expression are intriguing because the amount and intensity of coloration on male anal fins can change over an individual’s lifetime in response to social conditions and lighting environment (Fuller et al., 2022; Fuller & Travis, 2004; Johnson & Fuller, 2015). These non-visual opsins may represent an additional mechanism through which fin coloration is regulated.

Finally, we examined population-level patterns in gene expression associated with habitat variation. Epigenetic mechanisms, including DNA methylation and histone acetylation, are known to contribute to the regulation of phenotypically plastic traits (Lafuente & Beldade, 2019; Laine et al., 2023), and genes involved in these processes were significantly upregulated in the swamp population, potentially affecting genomic regions associated with blue coloration plasticity. The swamp population used in this study occurs just downstream from Lake Okeechobee within the sugar canal system of south Florida, where water is likely enriched for agricultural chemicals. Consistent with habitat-related influences, genes enriched for GO terms associated with ion and heme binding and responses to external stimuli were upregulated in the swamp population relative to the spring population. Our findings parallel work by Whitehead and colleagues on population-level transcriptomic divergence associated with chronic chemical exposure in the mummichog (*Fundulus heteroclitus*) (Crawford et al., 2020; Whitehead et al., 2011). Studies of pollution-tolerant mummichog populations have demonstrated that long-term exposure to contaminated habitats leads to consistent, population-specific shifts in gene expression, often involving pathways related to stress response and the cytochrome P450 family (Reid & Whitehead, 2016; Whitehead, 2013; Whitehead et al., 2010).

The extraordinary diversity of anal fin coloration in bluefin killifish illustrates how sexual dimorphism, polymorphism, and phenotypic plasticity can arise from overlapping genomic architectures. Our results suggest that sexual dichromatism is built upon broadly conserved pigmentation pathways that remain transcriptionally active in adult males, consistent with the dynamic and socially mediated nature of melanin-based signaling. The absence of strong sex differences in androgen receptor expression points to endocrine regulation acting through circulating hormone levels or antagonistic pathways involving progesterone receptors. Male color polymorphism is superimposed on this shared sexual framework in ways that reflect the mechanistic basis of coloration: structurally based blue coloration is associated with greater regulatory divergence and complexity, whereas pigment-based variation among non-blue males appears to involve more limited transcriptional differentiation. Together, these results support a model in which sexual dichromatism and male polymorphism are produced through a combination of shared sex-limited regulatory programs and morph-specific transcriptional variation, providing a foundation for testing how selection and environment jointly shape the evolution and expression of color traits.

## Author Contributions

R.K. and R.C.F. designed the study and collected the samples for RNA sequencing. C. Cheng extracted DNA for the genome and helped troubleshoot RNA extraction from minute fin materials. J. Catchen and G. Madrigal led the efforts on genome assembly and annotation and provided guidance on analyses of gene expression. R.K. performed laboratory work and data analysis of the transcriptomic data and drafted the manuscript. All authors contributed to manuscript revisions and approved the final version of the manuscript.

## Acknowledgements

We thank Elijah Davis and members of the Fuller lab for help with fish husbandry. This work was partly supported by the USDA National Institute of Food and Agriculture, Hatch project 1026333 (ILLU-875-984) through a Program in Ecology, Evolution and Conservation Biology summer research grant to RK. The genome assembly was supported by a University of Illinois Research Board grant to JC and RCF.

## Data Availability

The raw PacBio Hi-Fi reads (SRR40153233), Hi-C reads (SRR40157203) and 10X Illumina reads (SRR40159134) generated for the *Lucania goodei* assembly, along with the raw RNA-Sequencing reads (SRR40144438-SRR40144476), are available on NCBI under BioProject PRJNA1512036. The genome assembly will be made available on NCBI. An additional copy of the assembly, associated gene annotations, and code for analyses will be available on Dryad.

## Ethics Statement

This work was approved by the University of Illinois Institutional Animal Care and Use Committee (#23145).

## Supplementary methods

To identify the differentially expressed genes (DEGs) involved in sexual dichromatism, we fit a model with population and sex as the independent variables, each with two levels: spring and swamp, and male and female, respectively. To examine the DEGs between blue and non-blue males, we used only male data and fit a model with population and blue status as independent variables, each with two levels: spring and swamp, and blue and non-blue, respectively. Genes exhibiting significant differences in expression among groups were identified using the moderated F-test implemented in the limma package. Genes with a Benjamini–Hochberg-adjusted P < 0.05 were used to generate heatmaps of normalized expression values.

## Supplementary figures

**Figure S1.**
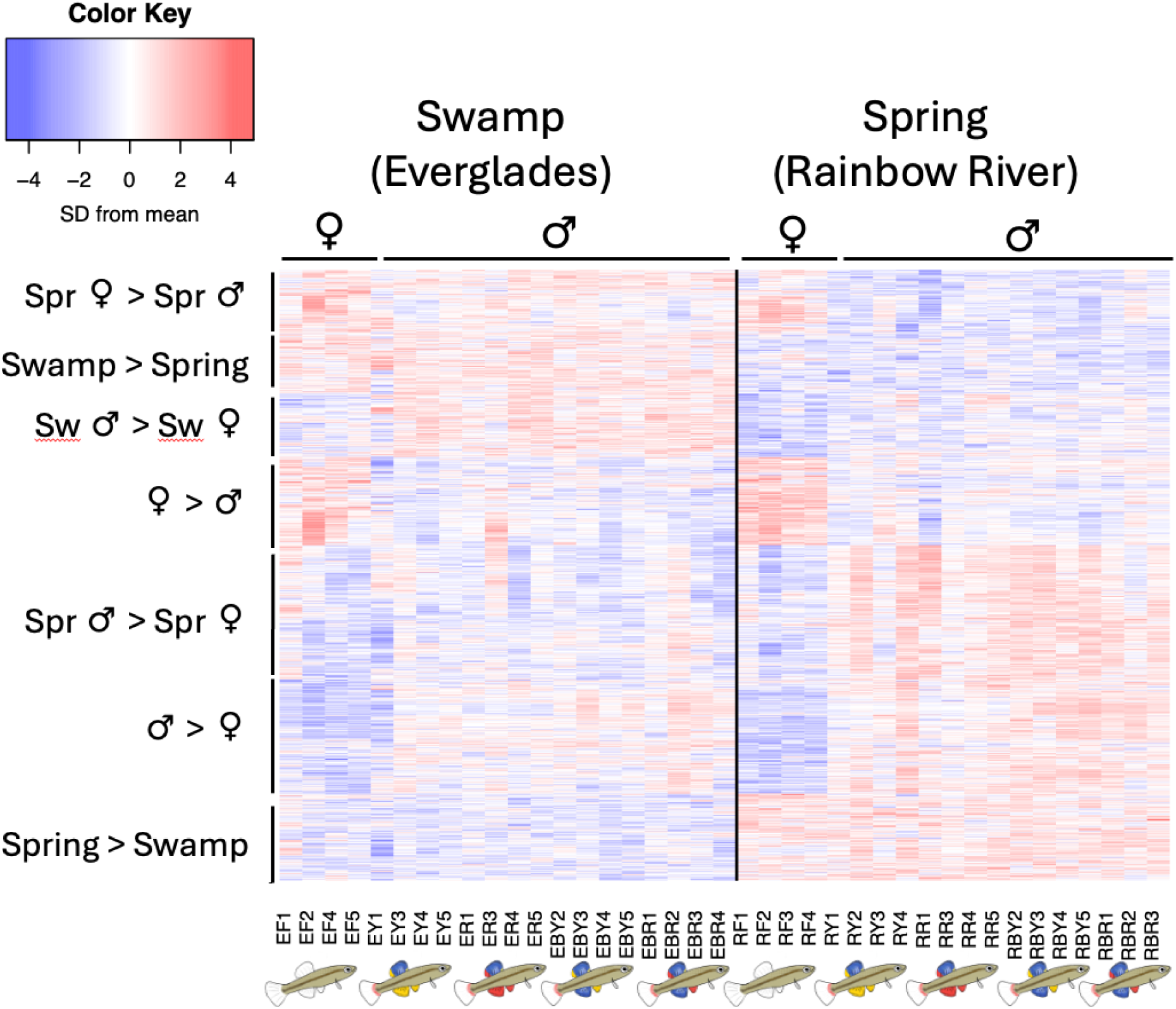
Heatmap with genes identified as differentially expressed genes in any comparison from the one-way ANOVA, showing the 1616 genes with differential expression between populations and sexes within each population. Each column is an individual; each row is a gene. Codes refer to the swamp population (Everglades, E) or the spring population (Rainbow River, R). F denotes females. R and Y denote red or yellow males. BR and BY refer to blue males with red or yellow pelvic fins, respectively.

**Figure S2.**
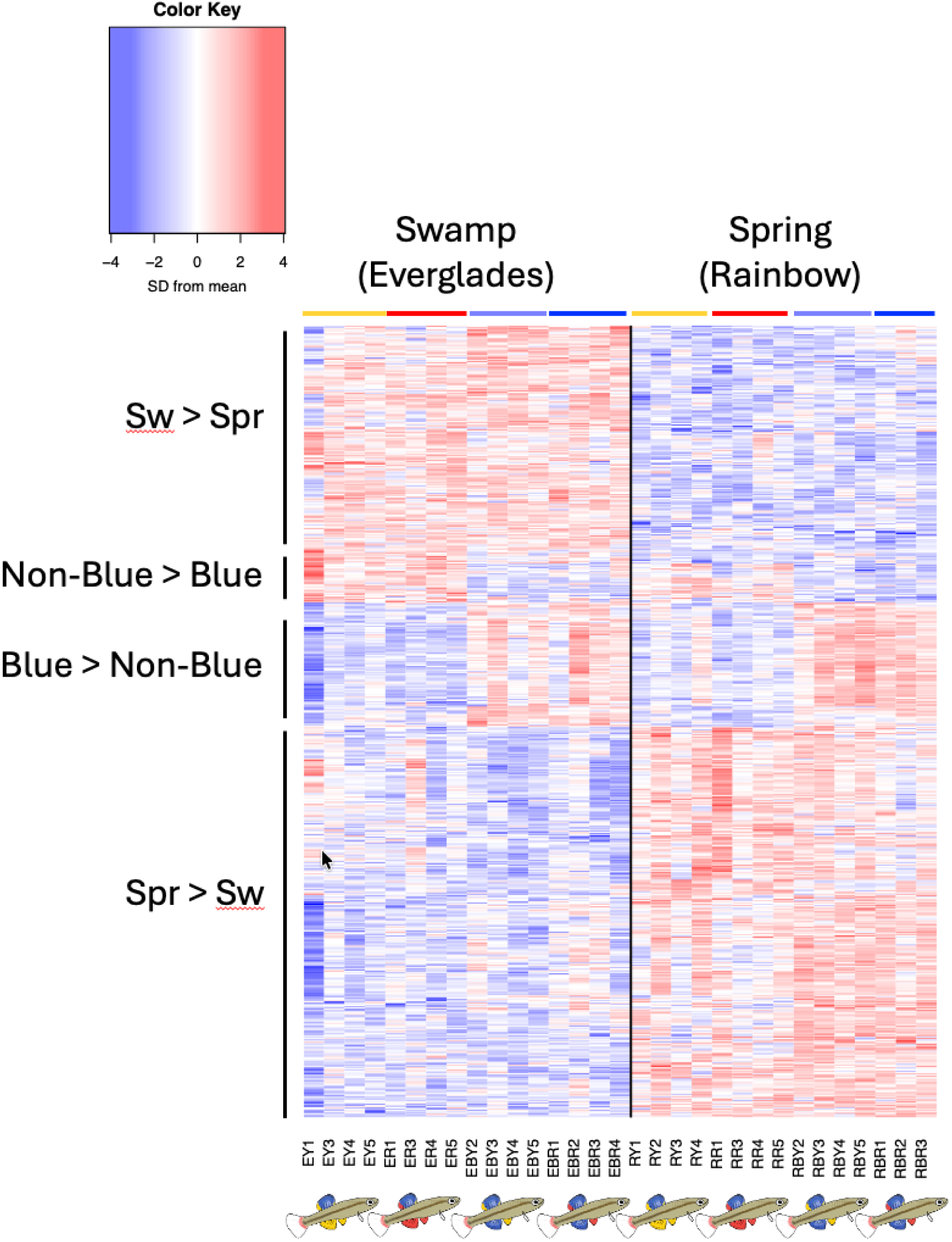
Heatmap with genes identified as differentially expressed genes in any comparison from the one-way ANOVA, showing the 557 genes with differential expression between populations and blue vs. non-blue males within populations. Colors above bars refer to yellow, red, blue carrying yellow (light blue), and blue carrying red (dark blue) males. Codes refer to the swamp population (Everglades, E) or the spring population (Rainbow River, R). F denotes females. R and Y denote red or yellow males. BR and BY refer to blue males with red or yellow pelvic fins, respectively.

**Figure S3.**
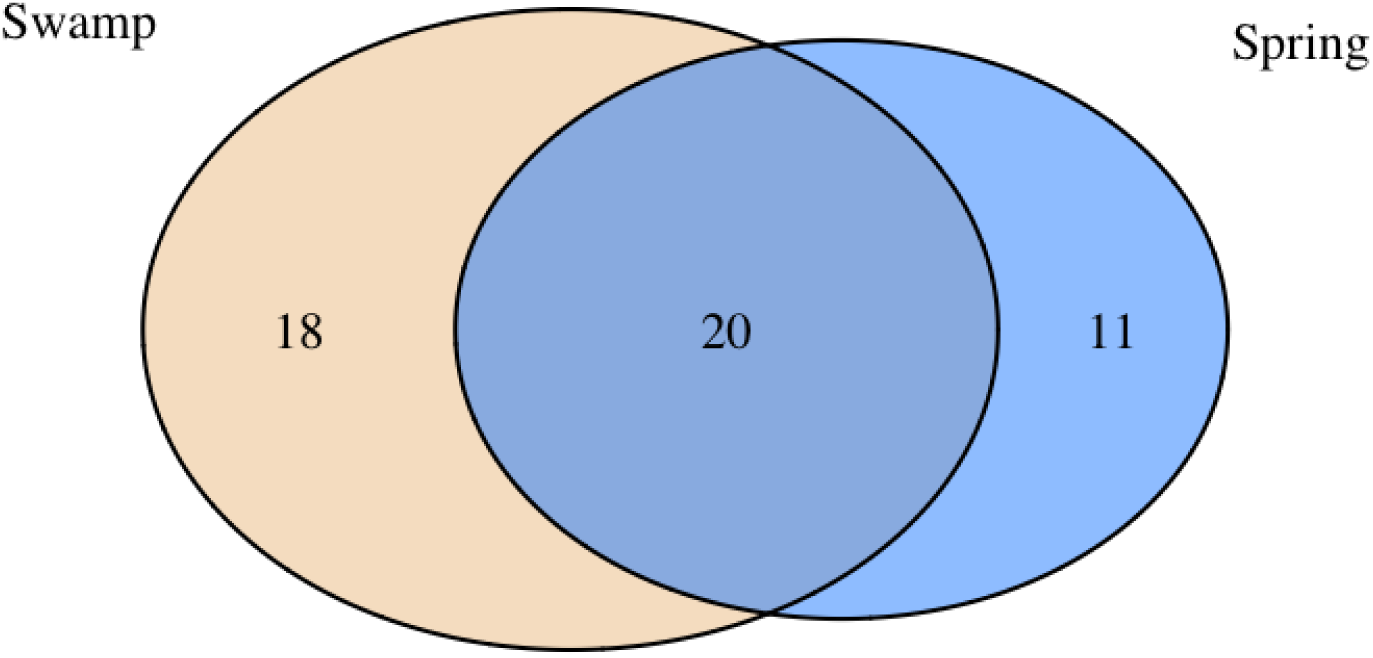
Number of significantly differentially expressed genes between blue and non-blue males from the two populations. Blue circle indicates the spring population, Rainbow river, and the yellow circle represents the swamp population, Everglades.

